# A model for collagen secretion by intercompartmental continuities

**DOI:** 10.1101/2023.09.21.558805

**Authors:** Louis Bunel, Lancelot Pincet, Vivek Malhotra, Ishier Raote, Frédéric Pincet

## Abstract

Newly synthesized secretory proteins are exported from endoplasmic reticulum (ER) at specialized subcompartments called exit sites (ERES). Cargoes like procollagen are too large for export by the standard COPII-coated vesicle of 60 nm average diameter. We have previously suggested that procollagen is transported from the ER to the next secretory organelle, the ERGIC, in TANGO1-dependent inter-organelle tunnels. Here, we show that intrinsically disordered domains of TANGO1 in the ER lumen generate an entropic contraction that pulls procollagen towards the ERES. Molecular gradients of pH and HSP47 between the ER and ERGIC generate a force in the range of tens of femtoNewtons (fN), which is sufficient to propel procollagen from the ER at a speed of ∼1 nm.s^-1^. This calculated speed and the quantities of collagen secreted are similar to its observed physiological secretion rate in fibroblasts, consistent with the proposal that ER export is the rate limiting step for procollagen secretion. Our theoretical model explains how cells can utilize molecular gradients to export procollagens at a rate commensurate with physiological needs.

**Significance Statement:** Procollagen cannot be exported from the endoplasmic reticulum (ER) by standard COPII-coated vesicle of 60 nm average diameter. We have previously suggested that collagen is transported from the ER to the next secretory organelle, the ERGIC, in TANGO1-dependent inter-organelle tunnels. ER and ERGIC differ in molecular composition including their pH and protein composition. We propose a mechanical/entropic ratchet model whereby molecular gradients of pH and the collagen chaperone HSP47, provide the energy to propel procollagen from the ER at a speed that matches the physiological rate of collagen secretion.

## Introduction

Secretory proteins are co-translationally inserted into the endoplasmic reticulum (ER), folded, and then collected at specialized domains called ER exit sites (ERES) for their transfer to the next station of the secretory pathway, the ER-Golgi Intermediate compartment (ERGIC). ERES are marked by COPII proteins, which are required for the export of secretory cargoes. COPII proteins assemble into a lattice on ER membranes to generate cargo-filled vesicles of 60 nm average diameter. Recent data suggest that COPII-dependent ER export can also happen through direct tunnels between ERES and the ERGIC, though the mechanistic details are rather scant (1, 2).

Secretory cargo capture into COPII-dependent vesicles is either through bulk flow or signal-mediated export. Bulk flow export occurs without signals, relying on passive cargo diffusion into an ERES. Signal-mediated export involves recognition of specific signals in cargo proteins by the COPII coat, leading to cargo concentration at ERES. Transmembrane proteins directly interact with Sec24 of the COPII inner coat, while soluble proteins or those without cytoplasmic signals require cargo receptors to facilitate an interaction with Sec24. Receptor and cargo exit the ER together. COPII protein assembly into a lattice or coat provides the motive force to concentrate cargoes into the export carrier for transport to the next secretory compartment (3, 4).

Compartmental identity changes gradually along the secretory pathway. There are differences in membrane lipid composition, soluble and membrane protein content, and ionic concentrations – including Ca^2+^ (5) and Zn^2+^ (6). Importantly, secretory organelle pH drops, from ∼7.4 in the ER to ∼6.5 at the ERGIC and cis-Golgi, to ∼5 in post-Golgi secretory granules (7). Stable gradients can also be maintained within micrometer long tubular organelles over time (8). It is not known if this electrochemical proton gradient contributes to the concentration and flow of secretory cargoes out of the ER.

A procollagen molecule has an extended, rigid, trimeric (triple-helical) region, which can reach 300 nm in length in the ER lumen, *i*.*e*. too long to fit in a COPII vesicle (9, 10). While it is clear that COPII proteins are required for procollagen export, no dedicated conventional cargo receptors have been identified yet. Procollagens are recruited to ERES by the ERES-resident protein TANGO1, which links procollagens to the COPII inner coat protein Sec23A. However, TANGO1 is unique among cargo receptor-like proteins in that it does not leave the ER along with the cargo – instead collagen moves forward, while TANGO1 is retained at the ERES. It remains unclear how procollagens are exported from the ER and whether factors other than cargo-receptor-COPII lattice assembly could provide a motive force.

An informative set of data that casts light on these questions comes from a final step in procollagen folding prior to its export from the ER. Once formed, the triple helical structure is unstable at ∼37°C. To stabilize collagen at these physiological temperatures, triple helical regions are held together by ‘clamps’ formed by Heat Shock Protein 47 (HSP47) dimers that can bind at several sites along the procollagen triple helix (11, 12). The binding of HSP47 to collagen exhibits exquisite pH sensitivity, almost completely bound at the near-neutral pH found in the ER (∼7.4) and almost completely dissociated by the lower pH found in the ERGIC (13). Live video microscopy of the ER-Golgi interface during procollagen secretion, at high spatiotemporal resolution, revealed that HSP47 dissociates from procollagen at the ERES and is retained at the ER (14).

In the lumen of the ER, HSP47 binds an SH3-like domain (SLD) of TANGO1 to recruit triple helical (fully folded) procollagen to the ERES. In addition to the SLD, the lumenal portion of TANGO1 (TANGO1_LUM_) comprises a large region of ∼1000 amino acids, most of which is predicted to be an intrinsically disordered region (TANGO1_LUM_ IDR). On the cytoplasmic side of the ERES, TANGO1 is composed of (Fig. 1A) two coiled-coil (CC) domains and a proline rich domain (PRD). Raote et al. showed that TANGO1 molecules can self-arrange into a ring at the ERES (15-17). Cytoplasmic domains also facilitate the formation of a tunnel between the ER and the ERGIC.

**Figure 1.**
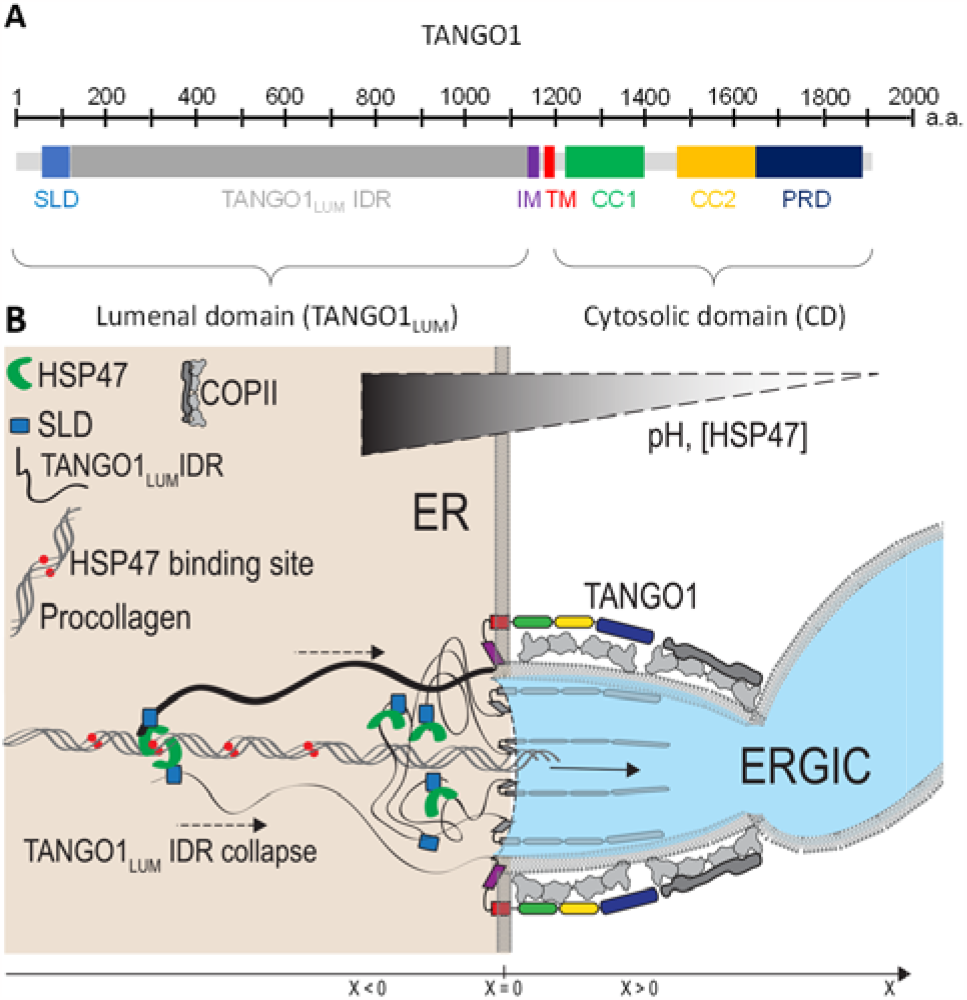
Model of procollagen transport by TANGO1. A. TANGO1 is a long protein that contains an ∼1000 r**e**sidue long intrinsically disordered TANGO1LUM IDR in the ER lumen, an intra-membrane domain (IM) followed by a transmembrane domain (TM) and two coiled-coil regions on the cytosolic side (CC1 and CC2). An SH3-like domain (SLD) is located at the N-terminus. B. The C-terminal coiled-coil regions of TANGO1 (not shown here) were suggested to open a tunnel between the ER lumen and the ERGIC. The lumenal portion of TANGO1 could separate the lumenal content of the ER and the ERGIC, creating a gradient of HSP47 and pH between the ER lumen and the tunnel. Some TANGO1LUM SLDs can bind to HSP47, which in turn binds to triple helical procollagen-generating a mechanical link that exert a force on procollagen. Because **of** the molecular gradients, this force is asymmetrical and drives procollagen movement. HSP47-binding sites are l**o**cated per pair every ∼10 nm on the procollagen. All quantitie**s** are defined along the x-axis representing the center of the tunnel. The junction between the tunnel and the ER is **pl**aced at x=0; x<0 is in the lumen of the ER and x>0 is in t**he** tunnel towards the ERGIC. Gradients of pH and [HSP47] are maintained, possibly by a network of TANGO1LUM IDRs.

An unusual combination of an intramembrane helix and a transmembrane helix (IM-TM) isolate the ERES membrane from the rest of the ER, acting as a partial diffusion barrier for lipids between the fusing ERGIC and the bulk of the ER (18). TANGO1_LUM_ along with the recruited collagen cargo, would form a densely packed protein network, perhaps functioning as a partial physicochemical barrier between the ERES and the ERGIC. The starting point of our model is based on the idea that this barrier maintains molecular gradients. Specifically, such a partial barrier could stabilize electrochemical gradients of pH or HSP47 content. The molecular gradients favor the binding of TANGO1_LUM_ IDR to procollagen in the ER lumen compared to the ERGIC. Tethering TANGO1_LUM_ IDR to procollagen reduces the number of accessible conformations, lowering the ensemble’s conformational entropy, and generates an effective entropic force that pulls away from the point of tethering. This entropic collapse resembles what was observed with transiently unstructured protein domains that provide sufficient forces to bring vesicles together (19). Here, we show that the molecular gradients can induce a total entropic motive force for secretory cargo transfer between the organelles. This force is sufficient to export procollagen from the ER at a physiological rate, even in the absence of a cargo-receptor.

## Results

### Presentation of the model

We begin at a stage where a tunnel has formed between the ER and ERGIC with TANGO1_LUM_ located at its base, facing into the lumen of the tunnel (Fig. 1B-C). A procollagen trimer spans the tunnel from the ER lumen to the ERGIC lumen, aligned along the axis of the tunnel. Procollagen contains binding sites for the chaperone-like protein HSP47, which is found in the ER lumen. Two HSP47 binding sites are located approximately every 10 nm on the procollagen molecule (12, 20, 21). HSP47 can simultaneously bind the SLD at the N-terminal region of TANGO1 (22, 23). This dual binding of HSP47 ensures a mechanical coupling between procollagen and some TANGO1 proteins of the ring. The affinities, on-rates and off-rates of HSP47 to procollagen have previously been measured at various pH (13, 24). These pH-dependent kinetics have been proposed to be a binding switch, such that HSP47 binds to procollagen at the neutral pH in the ER and then unbinds as the pH drops in the early secretory pathway (12). There is a considerable change in HSP47 affinity for collagen observed at the pH values as seen in these secretory compartments, from 6 to 7.5. For instance, the off-rate of HSP47 is 0.192 s^-1^ at pH 6 and 0.028 s^-1^ at pH 7.5. The affinity and rate constants between HSP47 and SLD do not vary with pH. We use the previously measured affinity 0.26 μM (22) and a dissociation rate of 0.03 s^-1^.

The establishment and maintenance of specific molecular content in the ER and ERGIC, and their use by TANGO1_LUM_ at the base of the tunnel, is key in our study. Two main gradients would be relevant for procollagen transport between ER and ERGIC – gradients of pH, and HSP47. These gradients define a zone of collagen binding at ER/ERES, and reduced binding in the tunnel/ERGIC. We will show that the gradients are sufficient to generate polarized and transient TANGO1/procollagen links, which produce enough force to propel procollagen from the ERES towards the ERGIC. We will evaluate the impact of these two gradients on the anterograde movement of procollagen. In the main text, we only consider a stepwise variation of pH and HSP47, while results of continuous gradients are presented in the supplementary information. It is well established that pH continuously decreases from ∼7.4 in the ER to ∼6 at the trans Golgi (7). Here, we will consider that the pH in the ERGIC and cis Golgi is ∼6.5.

There are a total of ∼172,000 TANGO1 molecules per cell (25), and ∼300 ERES per cell (26), i.e., an average of 500-600 TANGO1 per ERES. Also, there are ∼8 million copies of HSP47 (25) in HeLa cells, located in the ER. Hence, at a first approximation, we estimate the concentration of HSP47 in the ER at ∼50 μM (see Materials and Methods).

Based on these data, we will study the exit of procollagen over a range of TANGO1 molecules per procollagen varying from 10 to 600 and a local **c**on-centration of HSP47 in the ER varying from 1 t**o** 500 μM.

### A single TANGO1 generates force to propel procollagen

The long TANGO1_LUM_ IDR at the base of the tunnel is largely disordered. Each IDR ‘thread’ can therefore be described as a polymer anchored at the membrane by the TANGO1 transmembrane domain and extending randomly on either side of the tunnel. The probability density of the position of the SLD depends on the persistence length,, typically 4 residues () and the number N of monomers, i.e. the number of repeats of this persistence length over the contour length of the TANGO1_LUM_ IDR, here N = 264. This probability density also varies with the accessible space. In a tunnel, the geometry resembles a one-dimensional space while a fully open space is in three dimensions. The expression of the probability density (per unit volume) along the axis of procollagen, x-axis, in a space of dimension d is:

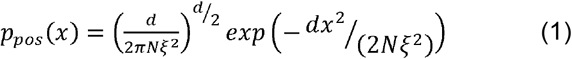

This symmetrical normal probability distribution is displayed in Fig. 2A for *d*= 3.

**Figure 2.**
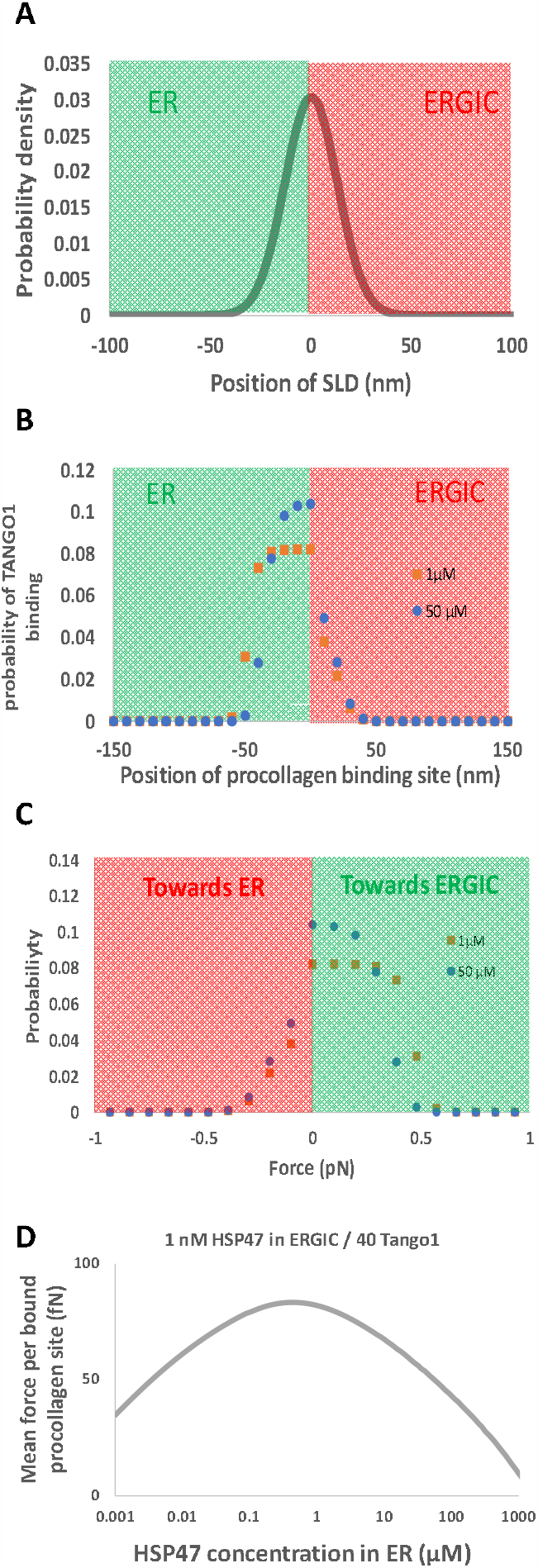
TANGO1 extension, binding probability and force. A. Because the TANGO1_LUM_ IDR is intrinsically disordered, the location of the SLD follows the symmetric gaussian distribution presented in Eq. 1. The case of a three-dimensional space is displayed here. B-C. Probability of the location (B) and force (C) of a TANGO1/procollagen link assuming 40 Tango1 molecules overall, a 1 nM HSP47 concentration, and pH 6.5 in the tunnel, and 1 μM (orange) or 50 μM (blue) and pH 7.4 in the ER lumen. Positive forces are towards the ERGIC and negative forces are towards the ER. These distributions are asymmetric as a result of the HSP47 and pH gradient. Regions shaded in green or red are in the ER or ERGIC, respectively in panels A, B and C. D. Mean force per bound procollagen site, obtained as the sum of the product of the force (panel A) and force probability (panel C) at each concentration for 40 Tango1.

Because of the pH and HSP47 gradients, the binding affinity of SLD to procollagen is different in the ER lumen and in the ERGIC. Convolving this difference and the SLD position distribution, provides the relative probability of a procollagen binding site to be occupied by an SLD (see Eq. 7 and associated text in the Materials and Methods). The examples presented in Fig. 2B display this asymmetric probability distribution and show limited variations with HSP47 concentration in the ER.

When a TANGO1_LUM_ stretches, it will try to retract, so that it can return to the relaxed conformation preferred by TANGO1_LUM_ IDR. If the TANGO1_LUM_ binds to a procollagen when it is stretched, its tendency to retract will generate a force that pulls procollagen towards the base of the tunnel. TANGO1_LUM_ will never stretch beyond 150 nm (probability < 10^−12^). Over this range, the extension of the TANGO1_LUM_ is equally modeled as a worm-like chain (WLC, (27)) or a freely-jointed chain (FJC (28)). Both models (WLC or FJC) predict the same pulling force, within a 1% margin. The force probability displayed in Fig. 2C is obtained by combining the position probability in Fig. 2B and the force associated with each position. The asymmetry of the curves in Fig. 2C shows that the resulting force tends to pull procollagen towards the ERGIC. The mean force per procollagen occupied by an SLD is presented in Fig. 2D, for HSP47 concentrations between 0.1 nM and 1 mM assuming 1 nM HSP47 in the tunnel on the ERGIC side and 40 Tango1. There is a change in direction of the asymmetry, i.e., the resulting force propels procollagen towards the ER, when HSP47 concentrations is in the mM range, which is non-physiological. This shift in direction occurs because all the collagen binding sites and SLDs are bound to HSP47 in the ER, precluding the formation of a bond between procollagen and Tango1. This large range of HSP47 concentrations over which the force favors a movement towards the ERGIC shows the versatility of the system – it is insensitive to large fluctuations in concentrations.

We now evaluate how several TANGO1 can collectively propel procollagen and induce procollagen movement. We present two approaches. In the first one, we assume all TANGO1 are at equilibrium. We studied the collective force generated by up to 600 TANGO1 per procollagen. The second approach presents simulations in which procollagen is completely unconstrained and movement occurs out of equilibrium. Because of the long simulation time, we could not go beyond 40 TANGO1 molecules per procollagen. Both approaches are complementary as they represent opposite situations (equilibrium vs. out of equilibrium) and provide similar quantitative result, showing the confidence in the conclusion.

### Force induced by several TANGO1 at equilibrium

To understand how movement can be triggered with several TANGO1 acting in parallel, we start with a procollagen molecule extending from the ER into the tunnel. The model considers procollagen and TANGO1 are at equilibrium. Under these conditions, all TANGO1 molecules anchored at the ring can randomly bind to procollagen via HSP47, as described in the previous section. The probability of how many TANGO1 molecules are bound to specific procollagen HSP47-binding sites can be calculated. Each given configuration of procollagen interacting with some molecules of TANGO1, is associated with a certain force experienced by procollagen. The mean force can be computed by summing over all such possible configurations (Fig. 3A, see Materials and Methods for detailed calculation). As in the case of a single TANGO1, because of the pH and HSP47 gradients, the net mean force is positive and propels procollagen into the tunnel. Because the contributions of all TANGO1 are additive, this mean force is larger than that produced by a single TANGO1 and can reach up to 2 pN.

**Figure 3.**
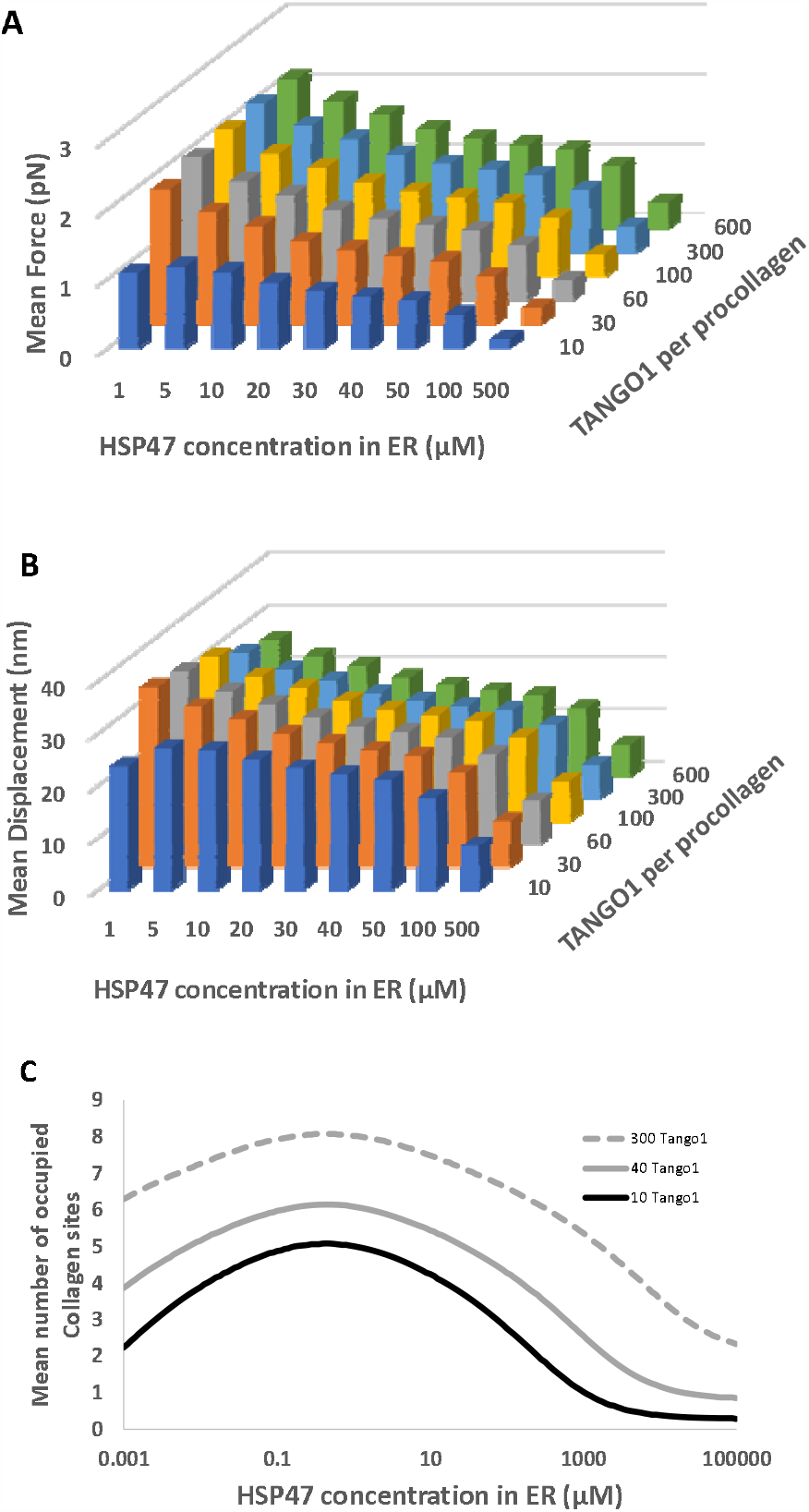
Equilibrium: force, displacement, occupied sites. A. Mean force on procollagen when several TANGO1 (10 to 40, right axis) are acting at equilibrium and for HSP47 concentrations varying from 1 μM to 500 μM in the ER lumen (front axis). B. Mean procollagen displacement (step) to dissipate the force exerted by TANGO1 on the procollagen. HSP47 concentration in the ERGIC is set at 1 nM. C. The number of occupied sites, i.e., bound to SLD via HSP47, depends on the total number of Tango1 per procollagen and the concentration of HSP47 in bulk. Three cases are presented here, 10, 40 and 300 Tango1, displayed in black, grey and dashed grey lines respectively.

To release the force, procollagen moves to a new position where there is no net force. The associated step size depends on the specific configuration of how many TANGO1 are interacting with procollagen at that moment. The mean step, presented in Fig. 3B for various HSP47 concentrations and numbers of TANGO1-procollagen interactions, remains in the range 10 - 30 nm (see Materials and Methods for detailed calculation). The procollagen displacement time can be estimated from the force it experiences and the equivalent viscosity of the medium (see Materials and Methods). This time is extremely short, less than 100 μs. Hence, when procollagen is unconstrained, it is unlikely that there is no time to reach equilibrium because procollagen moves too quickly compared to the kinetics of Tango binding and unbinding. Equilibrium can be reached if procollagen movement is retarded, *e*.*g*., by locally high viscosity, or transiently clamped by interactions with the membrane opposing the movement or by slow association of the procollagen trimer. In the latter case, movement will only occur when the clamp is released (growth of the tunnel or formation of the trimer) in the two examples), procollagen moves with a speed that varies linearly with the frequency of clamp release. This frequency must be slower than or commensurate with that of TANGO1 detaching from procollagen which is approximately the product of the off-rate of HSP47, 0.1 s^-1^, and the number of HSP47-procollagen interactions (below 10, see Fig. 3C). Hence the speed would be less than 20 nm.s^-1^.

### Movement of unconstrained procollagen

To obtain quantitative information on the movement of unconstrained out-of-equilibrium procollagen, the best approach is to perform simulations. We chose to start with a procollagen that is initially positioned at the base of the tunnel. TANGO1 molecules can bind randomly to procollagen, inducing a force thereby generating a step-wise movement. Thus, the first step occurs when the first TANGO1 binds. The binding and the subsequent movement bring this site of TANGO1/procollagen interaction to the base of the tunnel where the net force on procollagen is zero. The length of the second step, which occurs when a second TANGO1 binds, depends on the distance between the two TANGO1/procollagen bonds (see Fig. 4A for description). The direction and amplitude of the subsequent steps depend on the type of event (TANGO1 binding or unbinding), and location (ER or ERGIC lumen). Examples are shown of traces that represent a single run of simulation showing fiber displacement with time (Fig. 4B) and their means (Fig. 4C). Because of the increase of computation time with the number of TANGO1, we limited the simulations to a maximum of 40 TANGO1 per procollagen.

**Figure 4.**
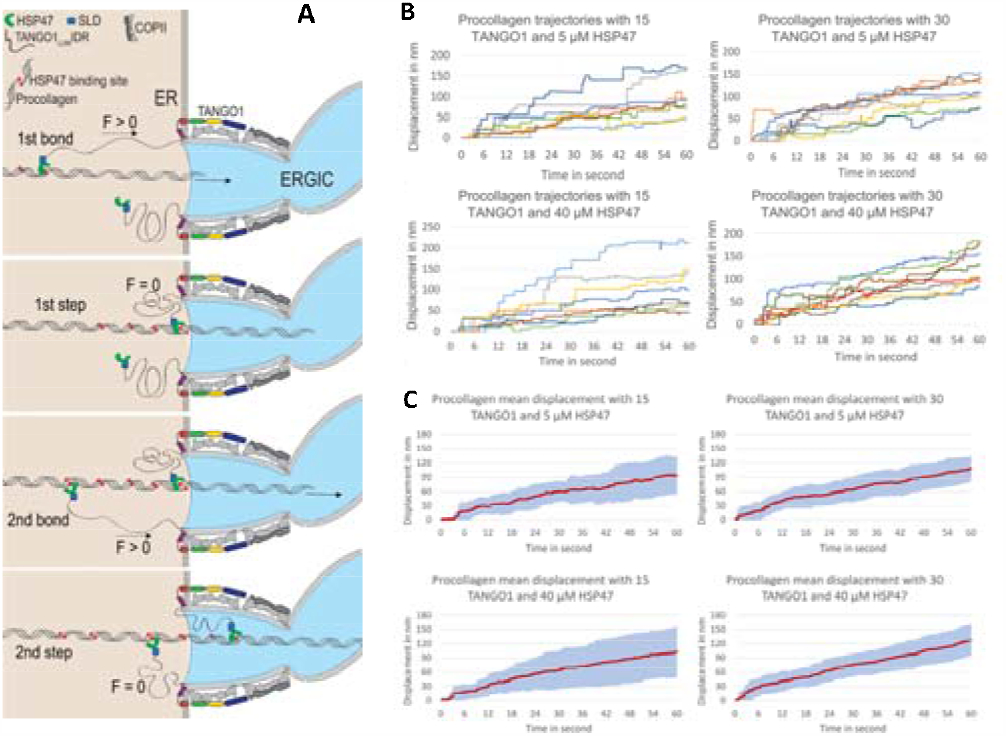
Movement of unconstrained procollagen. A. Schematic of the first events leading to procollagen movement. The first TANGO1 binds to procollagen (top panel), it applies a force that propels the procollagen until the force disappears (second panel, the arrow indicates the length of the step). A second TANGO1 binds (third panel) leading to a second step (bottom panel) and the cycles continue by attachment and detachment of TANGO1. B. Examples of trajectories (displacement of procollagen) in time for 8 simulations per conditions. C. Mean of the displacements over 15 trajectories. The blue shaded area corresponds to the standard deviation. In A and B, left panels are for 15 TANGO1, right panels for 30 TANGO1, top for 5 μM HSP47 in the ER lumen and bottom panels for 40 μM HSP47 in the ER lumen.

Despite the fact that both forward and backward movements of the procollagen occur within the 60 seconds simulations, the general features of the movement forward are reproducible between the various traces Fig. 4C. There are three phases. The first phase is a lag phase, waiting for the first TANGO1/procollagen bond to form. It lasts for a 3-4 seconds, when there are only 15 TANGO1 (Fig. 5C). The lag phase reduces as the number of TANGO1 molecules is increased. Fewer than 40 TANGO1 molecules are sufficient to make this lag phase disappear (Fig. 4B).

**Figure 5.**
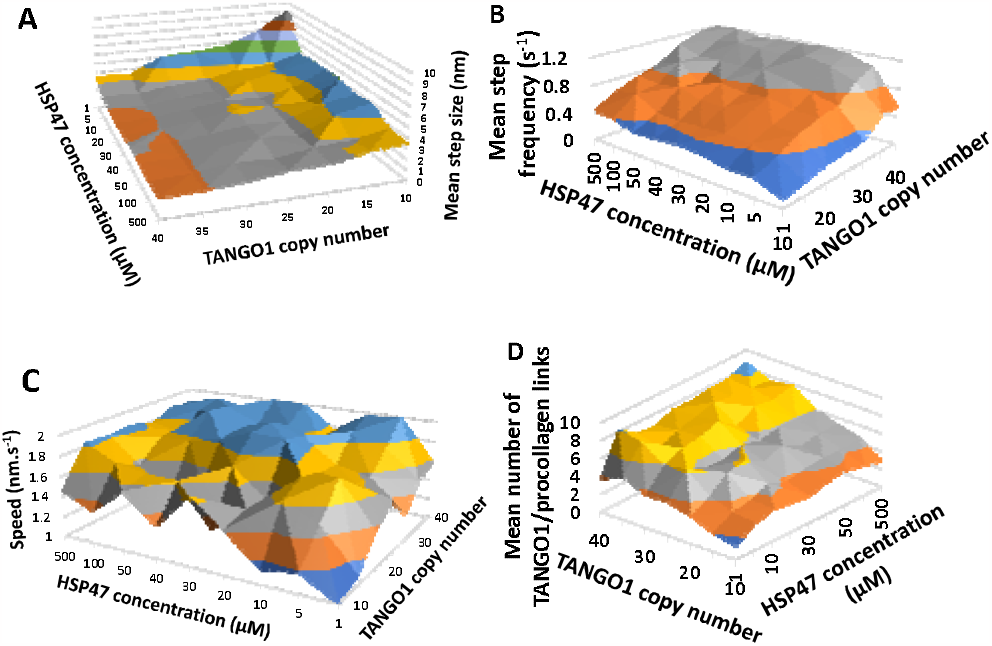
Physical characteristics of the trajectories. A, B, C and D display respectively the mean step size, step frequency, speed and TANGO1/procollagen links for 15 to 40 TANGO1 copy numbers and 1 to 500 μM HSP47 in the ER lumen.

During the second phase, only one or two TANGO1 are bound to procollagen. These bonds are almost exclusively in the ER. They need to be moved to the base of the tunnel or inside the ERGIC, *i*.*e*. over a large distance to dissipate the force. The resulting steps are large, ∼ 10 nm and the movement is fast, 10 to 20 nm.s^-1^, see for instance the fast movement between 2 and 3 s for 15 TANGO1 (Fig. 4C).

During the third phase, a steady-state is reached and procollagen advances at a uniform speed. The step size and their frequency depend on the number of TANGO1 molecules and the HSP47 concentration. Their variations are displayed in Fig. 5. Varying the number of TANGO1 or HSP47 concentration significantly changes the step size and frequency, between 1 and 10 nm and 0.1 and 1.2 s^-1^, respectively. However, these variations in TANGO1 numbers and HSP47 concentration almost cancel each other out, making the speed of procollagen movement almost uniform in the ranges investigated here, between 1 and 2 nm.s^-1^, with a maximum observed for 40 TANGO1 and 40μM unbound HSP47.

## Discussion

### Energetic principle underlying the model

Our model suggests that collagen export from the ER is carried out by a mechanochemical ratchet brought about by the entropic contraction of TANGO1_LUM_ IDRs surrounding the ERES. When the disordered region is partially extended it increases the probability of the SLD capturing a trimeric procollagen in the ER lumen at a distance away from the ERES. As one end of the TANGO1_LUM_ IDR is connected to the TANGO1 transmembrane and therefore attached to the ER membrane, this reduces the ensemble’s conformational entropy, generating an effective entropic force that pulls the SLD toward the ER membrane and propels procollagen from the ER to the ERGIC. The model, based on this principle, is summarized in Fig. 6. HSP47 binds procollagen and TANGO1 in the ER lumen and dissociates in the tunnel connecting them. TANGO1_LUM_ IDR that stretch in the ER lumen and are attached to procollagen exert a force that propels procollagen towards the ERGIC. Molecular gradients, *e*.*g*., pH and/or HSP47 concentration, are required to polarize the movement (Fig. 6). Hence, at least one of these gradients must be maintained for sustainable procollagen transport. This is actively achieved by mechanisms that recycle HSP47 towards the ER lumen and keep a constant pH in the ER and in the ERGIC. These mechanisms are an indirect energy source of procollagen movement.

**Figure 6.**
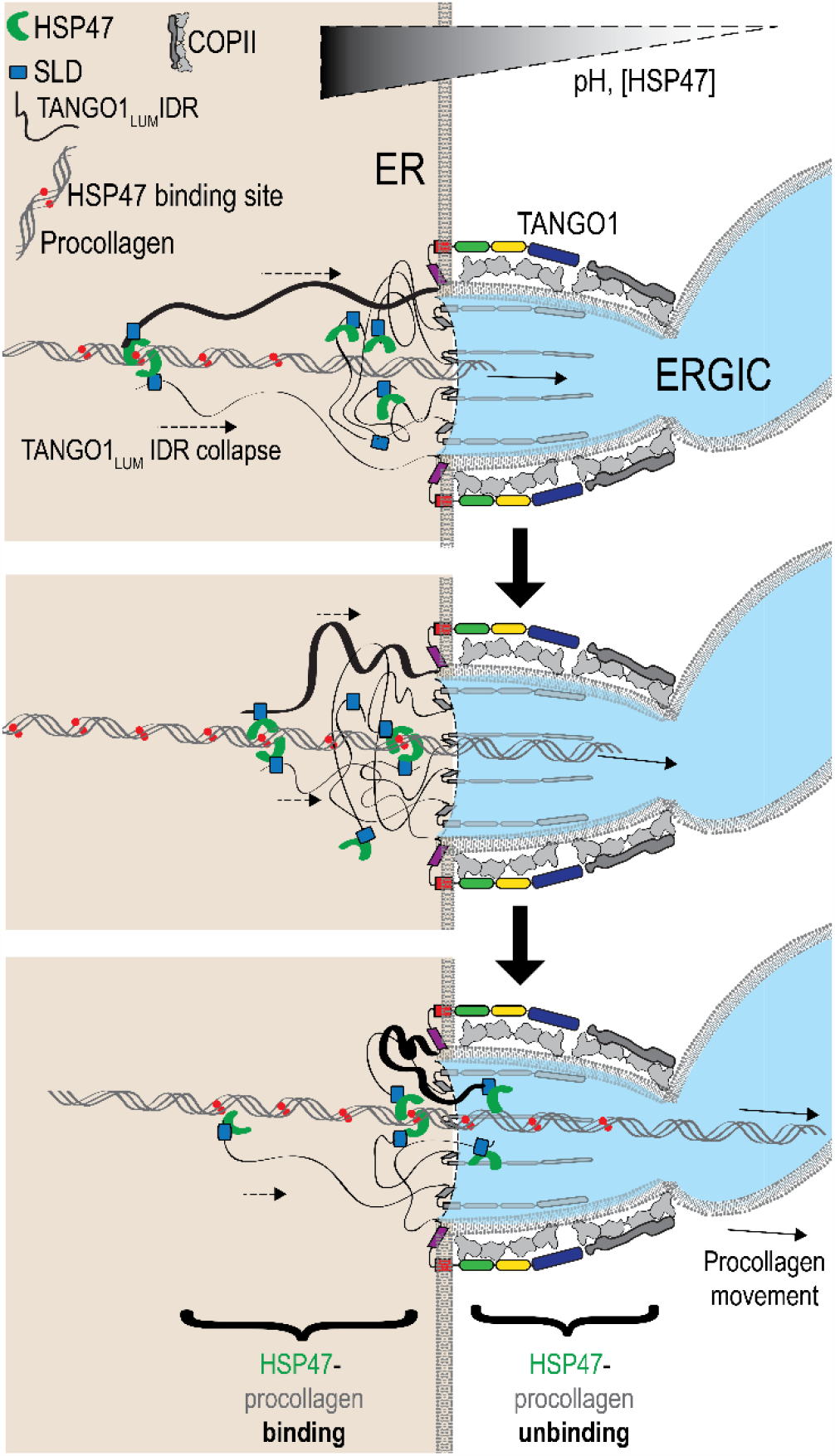
Diagram depicting how procollagen movement is achieved

### Stability of the system

Secretory traffic itself is pH-dependent, an effect that has been attributed to pH-sensitivity of processes and proteins involved in membrane fusion or fission. Here we show that a pH gradient between two compartments in the early secretory pathway could produce sufficient motive force to export cargoes from the ER without a need for a bona fide cargo receptor to drive cargo concentration.

Both the equilibrium and unconstrained models predict similar speed of procollagen transport, ∼1-10 nm.s^-1^. The wide range of HSP47 concentrations and TANGO1 copy number over which the speed remains constant shows the flexibility and stability of the system. ERES do not need to always contain exactly the same number of TANGO1 molecules per procollagen and the HSP47 concentration can vary over several orders of magnitude without affecting the efficiency of procollagen transport. The mean number of TANGO1/procollagen links varies (Fig. 5D) but always remains between 10 and 30% of the total number of TANGO1 per procollagen. On the ER side, only a small number of TANGO1 are bound to procollagen and are sufficient to produce significant movement. When there are more links between TANGO1 and procollagen, the movement is smoother but the speed remains the same. Typically, 70-90% of the TANGO1_LUM_ are not bound to procollagen and can maintain any other inter-TANGO1 cohesion.

The results presented here are based on the assumption that there is a stepwise variation of both HSP47 and pH at the base of the tunnel because of the TANGO1_LUM_ that can physically separate the ER lumen and the tunnel/ERGIC content. If only one of these two gradients actually exists, the conclusions remain qualitatively, as shown in Fig. 2D when the HSP47 concentration is the same in the ER and in the tunnel (data point for 1nM of HSP47in the ER). Similarly, a situation of a slow gradient of HSP47 concentration or pH would not affect the outcome significantly (see Supplementary information).

### Origin of molecular gradients at the ERES and experimental tests

In this study we have used pH and HSP47 as representative molecular gradients, but the outcome will hold true for any gradients that affect TANGO1-procollagen binding / unbinding in the ER and ERGIC. To study the putative origin of these gradients, HSP47 and protons are to be considered separately because they have very different properties: HSP47 is large, several nm in size, and diffuses slowly essentially following Stokes-Einstein equation for diffusion. Protons on the other hand are very small (∼0.1 nm) and diffuse in water according to the Grotthuss mechanism. In the following we will model the compartment that encompasses the tunnel and part of the ERGIC as a 500 nm long tubule with a diameter of 100 nm.

Regarding HSP47: it is easy to envision its distribution as a gradient. This type of protein gradient is observed in several systems including at the ERES. Such gradients of predominantly ER-resident proteins have already been described that are COPII-dependent (29) or cargo-dependent (30). It is also reasonable to consider that very high density of TANGO1_LUM_ IDR, cargo proteins, and cargo receptors, at the base of the tunnel creates a tight spaghet-ti-like network of disordered chains that may completely prevent HSP47 from entering the tunnel, or, at least significantly retard its diffusion towards the tunnel, thereby generating a gradient. It is worth noting that HSP47 bound to SLD and procollagen is brought in the tunnel by collagen. Because the affinity between HSP47 and SLD is ten times stronger than that between HSP47 and procollagen sites (260 nM vs. 2.25 μM), an HSP47 arriving in the ERGIC detaches primarily from procollagen. If it is also released from SLD, a single free HSP47 corresponds to a concentration of ∼500 nM in the tubule. Hence, this free HSP47 quickly rebinds to an SLD and is brought back to the ER by the spontaneous random movement of the corresponding TANGO1 molecule.

Regarding the proton gradient: the tubule contains ∼0.7 proton at pH 6.5. A freely diffusing proton would leave the tubule in approximately 10 μs, which is equivalent to a flow of 70,000 protons per second, which is 700 times the influx by as single V-ATPase (31). It is well-established that secretory pathway membranes are enriched in V-ATPase, Na/H exchange pumps, and other proteins that import protons into organelles. A lysosome of 250 nm average diameter is approximately of the same size as the tubule proposed here. Thus, it is unlikely that proton pumps will be sufficient to compensate for the protons freely exiting this tubule. However, pH gradients have been visualized in smaller tubular extensions in individual lysosomes (8). it is very possible that protons cannot freely diffuse out of the tunnel because the Grotthuss mechanism is based on local water organization that may easily be disrupted by the presence of the dense protein network of at the base of the tunnel. We assume that this impaired Grotthuss mechanism likely retards proton diffusion by several order of magnitude. For instance, if the diffusion is reduced 100 times, the influx of protons will be sufficient to maintain a pH on the ERGIC side of the tunnel. In terms of number of V-ATPase involved, it is well-established that each V-ATPase consumes ∼30 ATP.s^-1^ and 10 protons are pumped for 3 ATP (32, 33). Thus, a single V-ATPase can pump up to 100 protons per second, meaning 7 V-ATPase would be necessary on the ERGIC to maintain the pH, which is reasonable. This calculation does not include the action of other proteins such as sodium-proton exchange pumps that may even amplify the process.

In summary, a protein gradient has been demonstrated for other similar proteins, and realistic within this context. A proton gradient maintained by proton pumps and sodium-proton exchange pumps is plausible.

Even though the it is reasonable to consider molecular gradients maintained by dense protein networks of TANGO1_LUM_ IDR, cargo proteins, and cargo receptors, it has yet to be shown experimentally. Optical approaches seem the most likely to succeed as pH-sensitive variants of GFP have been directed to specific organelles (34). Sensitive measurements can be performed by using signal sequence-tagged GFP variants, including pHluorin (35) and dual-emission GFP (deGFP). More recent DNA-based sensors (36), such as CalipHluor (8) provide simultaneous quantitative information about both Ca^2+^ and pH, if they could be appropriately targeted to ERES The main difficulty is to visualize the tiny volume of the tubule at the time when it is assembled. High spatio-temporal resolution of the site using such reagents, could provide quantitative insights into a proton gradient.

### Comparison with physiological procollagen secretion

The predicted speed of a few nm.s^-1^ for transport of procollagen from the ER to the ERGIC can be compared to the physiological rate of procollagen secretion in cells. It was recently established that fibroblasts secrete ∼100,000 procollagen molecules per hour (37). Procollagen is roughly 300 nm long. Hence, fibroblasts secrete a quantity of approximately 30 mm procollagen per hour, i.e., 10,000 nm procollagen per second. There are about 180,000 TANGO1 per cell. Assuming, 60 TANGO1 propel procollagen any time translates to export of 3,000 procollagen molecules from the ER. This corresponds to a speed of ∼3. nm.s^-1^, which is consistent with the predicted value.

This model is based on the ER-ERGIC transport of procollagen in a vertebrate system, but represents a general mechanism where electrochemical gradients of ions and proteins can be used to provide the motive force to transfer components from one compartment to another.

### Building the ER/ERGIC tunnel

As a final comment, it is worth noting that the TANGO1-induced force presented here may facilitate the initial formation of the ER/ERGIC tunnel. Before the two compartments are connected, there is no pH or HSP47 gradient. However, TANGO1 will still bind to procollagen up to 60 nm away from the membrane. Since procollagen is exclusively in the ER, there is a “gradient” of procollagen binding sites and the global force generated will push on the membrane, starting a bud on the cytoplasmic face that may extend 60 nm towards the ERGIC to balance the pulling and pushing force. This deformation of the ER membrane can act synergistically with COPII proteins to build an export route.

Once the transport step is concluded and collagen has been transferred to the ERGIC, it is conceivable that ERES machinery disengage to separate the ERES and ERGIC continuity for a subsequent round of cargo transport. Alternatively, when there is no cargo to contribute to the ER-ERGIC diffusion barrier, then these local ionic gradients are abolished.

While the mechanism presented here relates to vertebrate TANGO1 / HSP47 – dependent procollagen export, the SLD could bind to other chaperones or cargo receptors that have similar pH-dependent activity profiles to utilize the same principle to export not only collagens, but also other secretory cargoes.

## Materials and Methods

### Procollagen modeling

Procollagen triple helices are rod-like complexes that are ∼300 nm long. In our model, a triple helical procollagen is transported through a tunnel generated by the function of TANGO1 from the ER to ERGIC. We have assumed that there is an infinite supply of procollagen. The movement of procollagen can thus be considered unidimensional. The origin of the movement axis, x= 0, is chosen at the position of the TANGO1 ring. Negative x values are located in the ER while positives are in the ERGIC. HSP47 binding sites are found in pairs, distributed evenly at 10nm intervals along the axis of procollagen. We represented this arrangement by modeling procollagen as a rigid long rod with dual HSP47 every 10 nm. The affinity, and on- and off-rates of HSP47 to procollagen binding have previously been measured and depend on the pH.

### HSP47 concentration in the ER

HSP47 is only located in the ER. There are ∼8 million copies of HSP47 per cell. Each cell has a volume of 2.5.10^3^ μm^3^ and the ER occupies 10% of the cell volume. Hence 8 million copies of HSP47 are randomly distributed in 2.5.10^2^ μm^3^ which corresponds to ∼50 μM.

### Binding procollagen and TANGO1

TANGO1 binds procollagen with low affinity (3.3 μM for procollagen IV (38)), which is neglected in our modelling. They can be indirectly connected by a shared HSP47 protein that is simultaneously attached to the SH3-like domain (SLD) of TANGO1 and one of the binding sites of procollagen. The probability that an SLD or a binding site on procollagen located at *x* is occupied by a HSP47 protein is respectively:

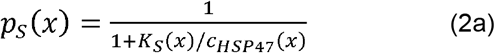

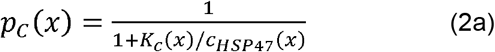

Where *K*_*S*_(*x*), *K*_*c*_(*x*) and *c*_HSP47_(*x*) are the affinities of HSP47 binding with SLD, with procollagen and the HSP47 concentration at *x* (as defined is Fig. 1).

### Force of TANGO1 on procollagen

When TANGO1 is connected to procollagen, it has to remain at that location, preventing its natural movement thereby generating a force pulling the procollagen towards the tunnel at *x* = 0. The value of this force, *f*(*x*), is very well characterized and can also be obtained using the WLC (27) or the FJC (28) model over the distance range we used here (between 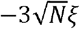 and 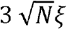).

### Procollagen movement modeling

Procollagen can be approximated as a cylinder of length 300 nm and diameter between 1.5 nm when HSP47-free and 8 nm when bound to HSP47. Following Broersma (39), we find that for a medium with the viscosity of water and a force of 50 fN, the initial speed of procollagen movement is between 10 and 60 μm/s. This speed vanishes exponentially with time during the displacement because of the fast decrease in force. The characteristic time for a 5 nm displacement is between 40 and 80 μs. As indicated in the main text, each step of the procollagen movement occurs with a frequency of 0.1 Hz or less, which shows that the displacement upon force can be considered as instantaneous. It could be argued that TANGO1_LUM_ locally increase the viscosity. However, this will impact the global movement only if the resulting viscosity is greater than 1,000 – 10,000 that of water.

### Equilibrium

In the equilibrium section, we assume that procollagen has already entered the tunnel and a pair of HSP47 binding sites is located at *x* = 0. Within the distance considered, there are 12 HSP47 binding sites in the ER and in the ERGIC. Hence, we consider 26 binding sites (13 pairs). Our goal here is to find the probability of all possible configurations in which some TANGO1 are connected to procollagen. Using the force corresponding to each configuration, we will obtain the mean force applied on procollagen. To compute all configurations, we assumed that TANGO1-SLD has full access to the whole space, *as shown in* Eq. (1) that is used in a three-dimensional system. In reality the dimensionality is probably lower, which will favor more bond formations, suggesting an underestimation of the force.

The local concentration of SLD along the axis is:

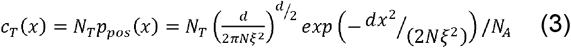

Where *N*_*A*_ is the Avogadro constant.

Hence the concentration of HSP47-bound SLD is:

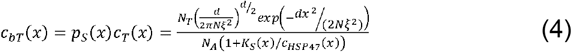

the probability of having a collagen binding site located at *x*_*i*_ is bound to Tango1 can be written as:

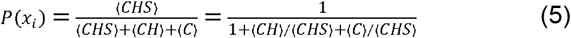

Where ⟨C⟩, ⟨CH⟩ and ⟨CHS⟩ are respectively the fractions of the time a collagen site is unbound, bound to HSP47 alone and bound to SLD via a shared HSP47. The ratios ⟨CH⟩ / ⟨CHS⟩ and ⟨C⟩ / ⟨CHS⟩ are equivalent to ratios of effective concentrations. Thus:

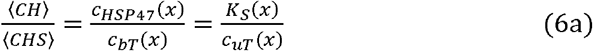

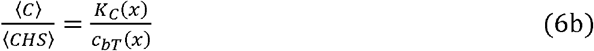

Where *C*_HsP47_(*x*) is the local concentration of HSP47 and *K*_*c*_(*x*) is the affinity between a collagen binding site and HSP47. The first equation assumes *K*_*c*_(*x*) is the same whether HSP47 is bound to SLD or not. The second equation is the equilibrium between C, CHS and bound SLD.

Hence, the resulting probability is:

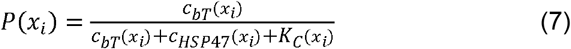

The mean force can now be expressed as:

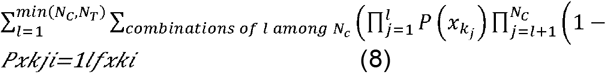

Where *l* is the number of occupied collagen sites in each combination 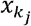 is the list of occupied sites and 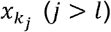 the unoccupied sites. *N*_*c*_ is the number of collagen sites (here, we set it to 26).

When a force is applied and freely moves, a displacement will occur to bring the force to zero. Noting 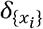 the displacement associated with configuration {*x*_*i*_}, we obtain the mean displacement per step:

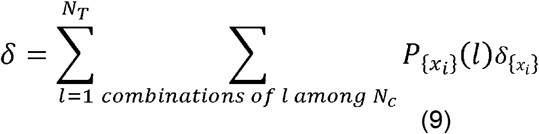

Where 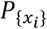 is the probability of combination {*X*_*i*_}.

### Out of equilibrium simulations

To study the displacement of procollagen in HSP47 and pH gradients, we used stochastic simulation algorithm considering the displacement of a single procollagen along the tunnel. At the start of each simulation, the procollagen is positioned at the entrance of the tunnel. HSP47 molecules are randomly bound to SLDs and procollagen sites according to the associated chemical reaction. The movement of pro-collagen is implemented in 10 ns steps. At each step, the SLD and HSP47 proteins freely diffuse (see Supplementary information), SLD/HSP47, HSP47/procollagen and SLD/HSP47/procollagen randomly bind or unbind and the procollagen moves by diffusion and under the force applied by TANGO1. Each simulation lasted an equivalent of 60 seconds. We tracked the position of procollagen extremity and of the force applied by TANGO1 on procollagen over the simulation time. Simulations were run for a number of TANGO1 in [10, 15, 20, 25, 30, 35, 40] and an external concentration of HSP47 in [1, 5, 10, 20, 30, 40, 50, 100, 500]. Fifteen trajectories were run for each pair of TANGO1 number and HSP47 concentration.

We assumed that binding of the reactants does not happen in any specific sequential order and that no cooperativity exists between the reactants, procollagen, HSP47 and TANGO1.

We also assumed that HSP47 in solution is constant over the simulation time, meaning that the simulated system is in contact with an infinite reservoir of HSP47. Free HSP47 molecules diffusing in the ER compartment are restricted to diffuse within the ER interval. Only HSP47 bound to procollagen or TANGO1 is allowed to cross the TANGO1 diffusion barrier. Whenever an HSP47 molecule in the ERGIC dissociates from both TANGO1 and procollagen, the HSP47 is removed from the ERGIC. Such immediate removal is abrupt, however live video microscopy, with high spatio-temporal resolution showed that HSP47 does not leave the ERES, suggesting that a removing and/or inactivation mechanism locally exists (14).

Details of the simulation procedure are presented in the Supplementary text.

The python codes for the Equilibrium and Out of equilibrium cases are provided in the Supplementary information.

## Acknowledgements

We thank Prof. Andreas Ernst and Prof. Felix Campelo for useful discussion. This work was supported by the LiquOrg synergy grant, ERC-2020-SyG-951146, awarded to V.M. and F.P. V.M. acknowledges the support of the Spanish Ministry of Science and Innovation to the EMBL partnership, the Centro de Excelencia Severo Ochoa and the CERCA Programme / Generalitat de Catalunya. I.R. acknowledges the support of Fondation pour la Recherche Médicale (grant AJE202210016216).

