## Supporting information for "A model for collagen secretion by intercompartmental continuities"

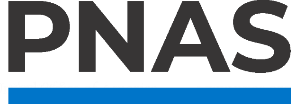

**Supporting Information for**

A model for collagen secretion by intercompartmental continuities

Louis Bunel, Lancelot Pincet, Ishier Raote, Vivek Malhotra, Frédéric Pincet

**This PDF file includes:**

Supporting text

Supporting references

Supporting Figures S1 to S2

Supporting Information Text

Simulation procedure

*Simulation strategy and chemical picture*

To study the displacement of procollagen in a pH gradient mediated by TANGO1 lumenal part, we used a stochastic simulation algorithm considering the displacement of a single procollagen in one dimension along the axis of the tunnel forming the link between the ERES and the ERGIC. We consider three chemical species namely: one 300 nm long procollagen molecule with thirty pairs of HSP47 binding sites evenly distributed every 10 nm, $N_{t}$ TANGO1_LUM_ IDR, each with a SLD and $N_{H}$ free HSP47 molecules, here $N_{H}=$50, see sub-step 3 below. Those species react in a selective manner forming *in fine* a SLD/HSP47/procollagen heterotrimer. Upon formation of the heterotrimer, TANGO1_LUM_ IDR applies a force on the procollagen, bringing it back towards the equilibrium position. We made the assumptions that binding of the reactants does not follow any sequence of events imposed by their chemistry and that no cooperativity exists between the reactants. Those assumptions are default hypothesis considering what is known from previous work in the field (1). Even though cooperativity for dual HSP47 binding site is suspected (2), it is not considered here because of the lack of precise kinetics and thermodynamics constants. We also assume that the concentration of free HSP47 in solution was constant over the simulation time, meaning that simulated system is in contact with an infinite reservoir of HSP47. Each time a free HSP47 binds to SLD or procollagen, another free one was added at a random position within the ER compartment to the system at the next simulation step. Similarly, whenever a HSP47 detaches from procollagen or SLD and becomes completely free, a free HSP47 chosen at random is removed from the ER compartment to maintain a constant number of free hsp47 molecules.

For this 1D simulation of procollagen export, we consider an axis along which the system evolved in time. As presented in the main text, the ER/tunnel junction is located at x=0 and is the location where TANGO1 IM/TM are assembled in a ring-like fashion. The ER compartment is delimited by the interval [-300 nm; 0] and the tunnel/ERGIC by x>0. We assume that procollagen fiber is already folded at t=0 and had the time to equilibrate with free HSP47 in solution. Thus we model the movement of a procollagen fiber pre-equilibrated with hSP47 and placed in contact at t=0 with an ERES containing a constant number of TANGO1_LUM_ IDR also pre-equilibrated with HSP47.

Initially, procollagen is positioned so its end is located at x=0, the $N_{H}$ free HSP47 are distributed randomly according to a uniform probability in the region x<0 and SLD ends of TANGO1_LUM_ IDR are distributed randomly according to the probability presented in Eq. 1 and Fig. 1A.

Then, at each time step, $\tau$=10 ns, we performed the following procedure with four sub-steps:

1. Displacement of the SLD at the end of each TANGO1_LUM_ IDR
2. Displacement of each free HSP47
3. Binding of all HSP47 in reactive range
4. Displacement of procollagen

As timescale of procollagen position relaxation is much faster than the unbinding characteristic time (ms *vs.* few seconds), a second slower time step $\tau_{reaction}$ is introduced during which unbinding can occur. The definition of $\tau_{reaction}$ is explained below (Eq. S.8) and its value is $\tau_{reaction}=26,43 \mu s$. Using this second timescale for unbinding shortens the simulation time without affecting the kinetics of the system_._

At each time step, $\tau_{reaction}$, we also performed the additional sub-step:

5. Random toss to test unbinding of each bond involving HSP47

Below are the descriptions of the 5 sub-steps.

*Sub-step 1: Displacement of the SLD domain at the end of each TANGO1LUM IDR.*

SLDs that are bound to procollagen through SLD/HSP47/procollagen trimers follow procollagen movement (see last sub step). For the other, free SLD and SLD carrying a HSP47 but unbound to procollagen, we used the description of TANGO1_LUM_ IDR presented in the main text. In brief, TANGO1_LUM_ IDR is composed of 1056 amino acids of length 0.35nm on average. Considering a persistence length of four amino acids, TANGO1_LUM_ IDR is modelled as a chain of 264 monomers of length 1.4nm. We modeled TANGO1 as a free-jointed chain which provides the value of the pulling force *vs*. the position of the SLD. The movement of SLD along the axis is driven by two contributions: random diffusion and movement produced by the pulling force. For the end-to-end diffusion coefficient, $D$, of the ideal chain modelling TANGO1, a value of 2.10^8^ nm²/s was chosen, following the work of Stirnemann et al.(3) on WLC modelling from experimental data on unstructured peptides. The diffusive movement of SLD at each time step, $\tau$=10 ns, follows a 1D diffusion step:

$x\left( t+\tau\right)=x\left( t \right)+\sqrt{2D\tau}$*ξ	(S.1)

With ξ a random number generated from a Gaussian distribution of parameter (0, 1) and $\tau$ the timestep. For the pulling force induced displacement over $\tau$, we used the force-extension curve presented in Fig. 1A and balanced it with a Stokes force, $\frac{k_{B}Tv}{D}$, to evaluate the speed $v$ of procollagen, where $k_{B}$ is the Boltzmann constant and $T$ the temperature. The corresponding displacement is $v\tau$.

By adding these two contributions, diffusion followed by force induced displacements over many $\tau$ on an unbound TANGO1_LUM_ IDR, we can reproduce the expected distribution of a random chain (Eq. 1 and Fig. 1A in the main text) which validates our approach.

*Sub-step 2: Displacement of free HSP47*

Considering the size of HSP47, we can approximate it as a sphere of 4nm radius. Using Stokes-Einstein relation, we can compute the diffusion coefficient of such an object to be 5.5.10^7^ nm²/s. In order to emulate the 1D diffusion of HSP47, we applied Eq. S.1 with the HSP47 computed diffusion coefficient.

Free HSP47 diffusing in the ER compartment are restricted to diffuse within the ER interval, x<0. Only HSP47 bound to procollagen or TANGO1 are allowed to cross the TANGO1 diffusion barrier. Whenever a HSP47 protein in the ERGIC compartment unbinds from both TANGO1 and procollagen, the HSP47 is removed from the ERGIC. Such immediate removal from the ERGIC may seem too rash, however HSP47 was found to be retained at the ERES (4) suggesting that a removing and/or retention mechanism exists at the ERES-ERGIC compartment junction.

*Sub-step 3: Binding of HSP47*

For HSP47 binding, we used the second order reaction approach presented in a 3D case by Andrews (5). Having two molecules diffusing in a defined volume and able to operate bimolecular reaction, it is possible to define a distance σ such that when two molecules get closer than the distance σ by diffusion reaction occurs. Such distance does not correspond to a real distance of reaction but is determined to recover the macroscopic kinetics observable experimentally. As stated before, our simulation take place along an axis with a segment [-300; 0] corresponding to the equivalent of the ER with a procollagen in it, composed of dual binding site every 10nm, so 60 binding sites exist in the ER simulation space. From Eq. S.3, we can express the number *x* of free procollagen binding sites considering n the number of total procollagen binding sites and the free HSP47 concentration noted [HSP47]:

$x= \frac{n*K_{D}}{\left[ HSP47 \right]+K_{D}}$	(S.2)

As $N_{H}$ the number of free HSP47 is kept constant during the simulation as if the system was connected to an infinite reservoir, *x* corresponds to the number of sites not bound to HSP47 at equilibrium. At steady state the number of procollagen binding sites undergoing association in a time step $\tau$ is the same as the one undergoing dissociation reaction. The number of dissociation reactions $r_{D}$ in a time interval $\tau$ corresponds to:

$r_{D}=m*k_{off}*\tau$	(S.3)

With *m* being the number of occupied sites ($m=n-x$*).*

The number of association reactions $r_{A}$ in a box of length *l* containing *n* binding sites and $N_{H}$ free HSP47 depend on the radius of binding σ as:

$r_{A}= \frac{2\sigma}{l}*N_{H}*\left( n-m \right)$	(S.4)

At steady state we have $r_{A}= r_{D}$ so a solution for σ can be found. Because the solution depends on *m* and consequently on *x*, we see that σ varied depending on free HSP47 concentration. As free HSP47 concentration increased, σ increased. This makes sense, considering that in this case the association reaction rate increased accordingly. Also, increasing the number of free HSP47 particles in the box, σ would decrease to give the same association overall despite this higher number of HSP47. As $r_{A}$ depends on the pair σ and $N_{H}$, we decided to fix the value of $N_{H}$ at 50 in order to have a good coverage of the simulation box but with an acceptable computation time. Such a value implies that on average we have a free HSP47 every 6nm. Considering the diffusion coefficient and the timestep $\tau$ of 10ns, at each timestep a HSP47 is moving by ~0.5nm on average meaning that on the timescale of the association/dissociation reactions, HSP47 distribution is homogeneous. With a timestep $\tau$ in the order of the tens of nanoseconds, σ was in the sub-nanometer range which is an unrealistic value for a radius of interaction one could expect. However, we are describing all particles of this system as point particles evolving over 1D meaning that they crossed each other in a systematic manner over time as they evolved on the same axis. This small value of σ incorporate the fact that not all collisions result in an association reaction.

*Sub-step 4: Displacement of procollagen*

The sum of all TANGO1_LUM_ IDR bound to procollagen through SLD/HSP47/procollagen trimers applies a force to procollagen. Modelling procollagen as a cylinder (see Materials and Methods in the main text), the speed of procollagen is obtained following Broersma (6) which directly gives the displacement during the simulation step $\tau$.

*Sub-step 5: Unbinding of HSP47*

As HSP47 is the chemical specie interacting with both TANGO1 and procollagen binding sites at each step of the reaction scheme (Eq. S2), we centered the chemical description of the system around HSP47. Two kinds of chemical reactions occurred in the present system: first order reactions of dissociation and second order reactions of heterodimerization. Those reactions were simulated by two different methods.

For the first order reactions of dissociation, we chose to describe the dissociation process considering the thermodynamics dissociation constant K_D_. In the general case of the dissociation reaction scheme of HSP47 with another specie S (either TANGO1 or procollagen binding site) we have:

*k_on_*

*k_off_*

$S-HSP47 S+HSP47$	(S.5)

K_D_ is described as:

$K_{D}= e^{\frac{-E^{\circ}}{kT}}= \frac{\left[ HSP47 \right][S]}{[S-HSP47]}$	(S.6)

From such a definition, the free S propensity f is given by:

$f= \frac{1}{1+ e^{\frac{E^{\circ}}{kT}}}$	(S.7)

Thus, the dissociation process of a pair S-HSP47 is computed using the procedure bellow at each time step of reaction $\tau_{reaction}$.

Coin = random value from uniform distribution in the interval [ 0; $1+e^{\frac{E^{\circ}}{kT}}$ ]

If Coin < 1: S-HSP47 heterodimer dissociates into S and HSP47, randomly separated by a distance of 1nm

Else: continue

This approach is valid at the condition that $\tau_{reaction}$checks the relation:

$k_{off}*\tau_{reaction}=f$	(S.8)

The values of K_D_ and k_off_ at the relevant pH are known (7, 8), hence, we can determine $\tau_{reaction}$. Using the measured values at pH = 7,5 we have K_D_ = 0,74µM and k_off_ = 0,028 s^-1^ in (7) we then obtain $\tau_{reaction}=26,43 \mu s$. Proceeding this way, we ensure to reproduce the thermodynamic equilibrium and the kinetic of the backward dissociation reaction corresponding to the values found experimentally by applying the Metropolis procedure at a well-defined frequency.

Alternate condition: continuous variation of the pH and HSP47 between the ER and ERGIC.

In the main text, we presented the case of a step-wise change of pH and HSP47 concentration at the base of the tunnel, $\boldsymbol{x=0}$. To show that the conclusions are not significantly altered by the initial conditions, we repeated the study assuming a pH that varies over 60 nm. The results are displayed in supplementary Figures S1 and S2 that can be directly compared to Figures 3 and 4 in the main text respectively. A situation in which both pH and HSP47 vary over 60 nm and only HSP47 varies over 60 nm are presented in Figures S3 and S4 respectively and can be compared with Figure 4. The magnitude of the forces and speed are not affected by the change in initial conditions.

Python code providing the average force of Tango1 on collagen at equilibrium, and mean collagen displacement to release this force.

```
"""
Created on Sat Sep  2 15:08:44 2023

@authors: Lancelot Pincet, Frédéric Pincet

Gives the average force of Tango1 on collagen at equilibrium, and mean collagen displacement to release this force.

Case with :
pH 7.5 in ER, 6.5 in Golgi
10nM HSP47 in ERGIC
pH gradient, HSP47 gradient
variable HSP47 in ER
variable number of Tango1 copies per collagen
"""

#imports
import itertools
import numpy as np
import time
import os
import matplotlib.pyplot as plt
import pandas as pd
import matplotlib
from numba import njit
from joblib import Parallel, delayed
from multiprocessing import cpu_count
from scipy.special import erf
import math

# =============================================================================
# User inputs

#files
path = r'' #Path where to save files

#Algorithm
do_parallel = True #True to loop on parallel values
ncores = cpu_count()-4 #Number of cores to use

#Loops
Ntango = [10,30,60,100,300,600] #Number of tango1 molecules
#Ntango = [30] #Number of tango1 molecules
HSP_ER = [1,5,10,20,30,40,50,100,500] #HSP47 concentration in ER [µM]

#Collagen definition
coll_max = 26 #maximum number of collagen sites
coll_min = 2 #minimum number of collagen sites
coll_d = 10 #Distance between two pairs of collagen sites [nm]

#Gradient on pH
pH_start = 0 #position start of gradient of pH [nm]
pH_end = 10 #position end of gradient of pH [nm]

#HSP47 concentrations
HSP_start = -30 #position start of gradient of HSP47 [nm]
HSP_end = 30 #position end of gradient of HSP47 [nm]
C_HSP2 = 1e-9 #HSP47 concentration in ERGIC [M]

#Tango1 polymer description
pers = 1.4 #persistence length in nm = 4 residues
Monom = 264 #nb of monomers = 1056 residues / 4 residues
Avogadro = 6.02e-1 #in molec.nm^-3.M^-1

#Affinities
KS = 0.26e-6 #Affinity of HSP47 with SH3 [M]
KC_ERGIC = 6.23e-6 #Affinity of HSP47 with collagen site in ERGIC [M]
KC_ER = 0.74e-6 #Affinity of HSP47 with collagen site in ER [M]


# User inputs
# =============================================================================

#Inializing collagen
poscoll = np.arange(2*coll_max) #id of collagen sites
xpos = np.floor((poscoll+1-coll_max)/2)*coll_d #Collagen pairs positions

#Initializing Tango1
cont = Monom * pers #contour length of Tango1 [nm]
sigma = np.sqrt(Monom*pers**2) #standard deviation
ppos = (3**(3/2)*np.exp(-3*xpos**2/2/sigma**2))/(2*np.pi*sigma**2)**(3/2) #density probability of SH3
alpha = abs(xpos/cont)
force = -np.sign(xpos)*4.14/pers*(1/(4*(1-alpha))-0.25+alpha-0.5164228*alpha**2-2.737418*alpha**3+16.07497*alpha**4-38.87607*alpha**5+39.49944*alpha**6-14.17718*alpha**7) #Force of polymer with end distance = alpha*contour_length

#Affinity of HSP47 with collagen site [M]
KC = np.empty(np.shape(xpos))
mask_pH1 = xpos < pH_start
mask_pH2 = (xpos >= pH_start) * (xpos <= pH_end)
mask_pH3 = xpos > pH_end
KC[mask_pH1] = KC_ER
KC[mask_pH2] = KC_ER + (KC_ERGIC - KC_ER)*(xpos[mask_pH2] - pH_start)/(pH_end-pH_start)
KC[mask_pH3] =KC_ERGIC

#Variable initializations
probaS = np.empty(np.shape(xpos)) #Probability that an SH3-like domain is bound to HSP47
probaC = np.empty(np.shape(xpos)) #Probability that a collagen binding-site is bound to HSP47

#combination calculations function
@njit()
def comb_calc(nc,Nc,comb,sites,p,p_tot,f,f_tot,Ptot,Nnorm,Ncproba,F,Fproba,Fabs,Fabsproba,disp,dispabs,dispproba,dispabsproba) :
    complem = np.asarray([i for i in sites if i not in comb]).astype(np.int32) #list of unbound sites
    P = np.prod(p[comb])*np.prod(1-p[complem]) #Calculating the probability of the combination
    Ptot += P
    Nnorm += 1
    Ncproba += nc*P
    Fint = np.sum(f[comb]) #force
    F += 1000*Fint
    Fproba += 1000*P*Fint
    Fabs += 1000*abs(Fint)
    Fabsproba += 1000*P*abs(Fint)
    if Fint != 0 :
        for shiftx in range(1,int(coll_max-Nc/2)):
            combadd = np.empty(len(comb)).astype(np.int32)
            combadd.fill(int((coll_max-Nc/2)+shiftx*np.sign(Fint))) #Need to go back to f_tot because some collagen sites may be out of the considered range
            Ftemp = np.sum(f_tot[comb+combadd])
            if np.sign(Ftemp) != np.sign(Fint):
                break
        disp += shiftx*10*abs(Fint)/(Fint-Ftemp) #Maybe not the best linear approximation but not too bad also
        dispabs += shiftx*10*Fint/(Fint-Ftemp)
        dispproba += P*shiftx*10*abs(Fint)/(Fint-Ftemp)
        dispabsproba += P*shiftx*10*Fint/(Fint-Ftemp)
    return Ptot,Nnorm,Ncproba,F,Fproba,Fabs,Fabsproba,disp,dispabs,dispproba,dispabsproba


# =============================================================================
# main

if __name__ == '__main__' : #Main script start

    #Starting code
    print('*****Script starting*****')
    tic = time.perf_counter()

    #Creating folder for gradient chosen
    prefix = '3D_'
    grad_path = path + prefix+ 'pH_{}_{}_HSP_{}_{}_nm/'.format(pH_start,pH_end,HSP_start,HSP_end)
    if not os.path.isdir(grad_path) : os.mkdir(grad_path)


    #Loops on Number of tango
    for Nt in Ntango :

        #Start Nt
        if do_parallel : end_string = ''
        else : end_string = '\n'
        print('Tango1 = {} molecules'.format(Nt),end=end_string)
        Nt_tic = time.perf_counter()

        #Making folder for number of Tango1
        tango_path = grad_path+'Tango1_'+str(Nt)+'/'
        if not os.path.isdir(tango_path) : os.mkdir(tango_path)


        #function to loop on HSPER values
        def HSP1_calc(HSP1) :

            #Start C_HSP1
            print('     HSP1 = {} µM'.format(HSP1))
            C_HSP1 = HSP1 * 1e-6 #HSP1 in µM C_HSP1 in M (ER)
            HSP2 = C_HSP2 * 1e+6 #HSP2 in µM C_HSP2 in M (ERGIC)
            HSP_tic = time.perf_counter()

            #Making folder for HSP47 concentration
            HSP_path = tango_path + 'HSP47_{}_{}_µM/'.format(HSP1,HSP2)
            if not os.path.isdir(HSP_path) : os.mkdir(HSP_path)

            #HSP47 concentrations
            C_HSP = np.empty(np.shape(xpos))
            mask_HSP1 = xpos < HSP_start
            mask_HSP2 = (xpos >= HSP_start) * (xpos <= HSP_end)
            mask_HSP3 = xpos > HSP_end
            C_HSP[mask_HSP1] = C_HSP1
            C_HSP[mask_HSP2] = C_HSP1 + (C_HSP2 - C_HSP1)*(xpos[mask_HSP2] - HSP_start)/(HSP_end-HSP_start)
            C_HSP[mask_HSP3] = C_HSP2

            #Probabilities
            probaS = 1/(1+KS/C_HSP)
            probaC = 1/(1+KC/C_HSP)
            CbT = probaS*Nt*ppos/Avogadro #local concentration of HSP47-bound SH3-like domains
            CuT = (1-probaS)*Nt*ppos/Avogadro #local concentration of HSP47-bound SH3-like domains
            p_tot = (1/(1+KC/CbT+KS/CuT)).astype(np.float32) #probability to have a collagen - Tango bond at each collagen site
            f_tot = force.astype(np.float32)

            #Initializing arrays for couple (NTango1,HSP47)
            Ptot_a,Fmean_a,Fabsmean_a,Ncproba_a,Fproba_a,Fabsproba_a,dispmean_a,dispabsmean_a,dispproba_a,dispabsproba_a = [],[],[],[],[],[],[],[],[],[],


            #Looping on number of collagen sites
            for Nc in range(coll_min,coll_max+1,2) : #number of collagen sites of interest
                print('          Collagen = {} sites'.format(Nc),end='')
                Nc_tic = time.perf_counter()
                sites = np.arange(Nc).astype(np.int32) #Defining the sites index for the permutations
                F,Fproba,Fabs,Fabsproba,Ptot,Ncproba,Nnorm,disp,dispabs,dispproba,dispabsproba = 0,0,0,0,0,0,0,0,0,0,0, #Reset before each Nc

                #Correcting P and f to match Nc
                p = p_tot[int(coll_max-Nc/2):int(coll_max+Nc/2)]
                f = f_tot[int(coll_max-Nc/2):int(coll_max+Nc/2)]

                #Calculating forces
                for nc in range(1,min(Nc,Nt)+1) : #For loop on the number of tango1 attached
                    combinations = itertools.combinations(sites,nc) #Calculates all combinations of nc elements in sites
                    for comb in combinations : #Looping on combinations
                        comb = np.asarray(comb).astype(np.int32)
                        Ptot,Nnorm,Ncproba,F,Fproba,Fabs,Fabsproba,disp,dispabs,dispproba,dispabsproba = comb_calc(nc,Nc,comb,sites,p,p_tot,f,f_tot,Ptot,Nnorm,Ncproba,F,Fproba,Fabs,Fabsproba,disp,dispabs,dispproba,dispabsproba)
                #Storing values
                Fmean = F/Nnorm
                Fabsmean = Fabs/Nnorm
                dispmean = disp/Nnorm
                dispabsmean = dispabs/Nnorm
                Ptot_a += [Ptot]
                Fmean_a += [Fmean]
                Ncproba_a += [Ncproba]
                Fproba_a += [Fproba]
                Fabsmean_a += [Fabsmean]
                Fabsproba_a += [Fabsproba]
                dispmean_a += [dispmean]
                dispabsmean_a += [dispabsmean]
                dispproba_a += [dispproba]
                dispabsproba_a += [dispabsproba]

                #End loops Nc
                Nc_toc = time.perf_counter()
                print(' took '+str(int((Nc_toc-Nc_tic)*100)/100)+' seconds')


            #End loops HSP
            HSP_toc = time.perf_counter()
            print('      took '+str(int((HSP_toc-HSP_tic)*100)/100)+' seconds')


            #Plotting
            saving_path = HSP_path
            matplotlib.use('Agg')  #Non interactive mode to refresh memory when savefig
            plt.figure()
            plt.plot(dispmean_a,'r')
            plt.title('Mean displacement')
            plt.xlabel('Number of bound collagen sites')
            plt.ylabel('dislacement [nm]')
            plt.grid(which='major', axis='both', linestyle='-',color='gray')
            plt.grid(which='minor', axis='both', linestyle='--',color='lightgray')
            plt.minorticks_on()
            plt.savefig(saving_path+'Mean diplacement.png', dpi=300)

            plt.figure()
            plt.plot(dispabsmean_a,'r')
            plt.title('Mean absolute displacement')
            plt.xlabel('Number of bound collagen sites')
            plt.ylabel('dislacement [nm]')
            plt.grid(which='major', axis='both', linestyle='-',color='gray')
            plt.grid(which='minor', axis='both', linestyle='--',color='lightgray')
            plt.minorticks_on()
            plt.savefig(saving_path+'Mean absolute diplacement.png', dpi=300)

            plt.figure()
            plt.plot(dispproba_a ,'r')
            plt.title('Displacement weighed by probability')
            plt.xlabel('Number of bound collagen sites')
            plt.ylabel('displacement [nm]')
            plt.grid(which='major', axis='both', linestyle='-',color='gray')
            plt.grid(which='minor', axis='both', linestyle='--',color='lightgray')
            plt.minorticks_on()
            plt.savefig(saving_path+'displacement_proba.png', dpi=300)

            plt.figure()
            plt.plot(dispabsproba_a,'r')
            plt.title('Absolute displacement weighed by probability')
            plt.xlabel('Number of bound collagen sites')
            plt.ylabel('displacement [nm]')
            plt.grid(which='major', axis='both', linestyle='-',color='gray')
            plt.grid(which='minor', axis='both', linestyle='--',color='lightgray')
            plt.minorticks_on()
            plt.savefig(saving_path+'absolute_displacement_proba.png', dpi=300)

            plt.figure()
            plt.plot(Fproba_a,'r')
            plt.title('Force weighed by probability')
            plt.xlabel('Number of bound collagen sites')
            plt.ylabel('Force [fN]')
            plt.grid(which='major', axis='both', linestyle='-',color='gray')
            plt.grid(which='minor', axis='both', linestyle='--',color='lightgray')
            plt.minorticks_on()
            plt.savefig(saving_path+'Force proba.png', dpi=300)

            plt.figure()
            plt.plot(Fmean_a,'r')
            plt.title('Mean Force')
            plt.xlabel('Number of bound collagen sites')
            plt.ylabel('Force [fN]')
            plt.grid(which='major', axis='both', linestyle='-',color='gray')
            plt.grid(which='minor', axis='both', linestyle='--',color='lightgray')
            plt.minorticks_on()
            plt.savefig(saving_path+'Mean Force.png', dpi=300)

            plt.figure()
            plt.plot(Fabsproba_a,'r')
            plt.title('Absolute Force weighed by probability')
            plt.xlabel('Number of bound collagen sites')
            plt.ylabel('Force [fN]')
            plt.grid(which='major', axis='both', linestyle='-',color='gray')
            plt.grid(which='minor', axis='both', linestyle='--',color='lightgray')
            plt.minorticks_on()
            plt.savefig(saving_path+'Absolute Force proba.png', dpi=300)

            plt.figure()
            plt.plot(Fabsmean_a,'r')
            plt.title('Absolute Mean Force')
            plt.xlabel('Number of bound collagen sites')
            plt.ylabel('Force [fN]')
            plt.grid(which='major', axis='both', linestyle='-',color='gray')
            plt.grid(which='minor', axis='both', linestyle='--',color='lightgray')
            plt.minorticks_on()
            plt.savefig(saving_path+'Absolute Mean Force.png', dpi=300)

            plt.close("all")
            matplotlib.use('QtAgg') #Putting back interactive mode

            #Saving data in excel
            df = pd.DataFrame()
            df['Ncproba'] = Ncproba_a
            df['proba'] = Ptot_a
            df['dispmean'] = dispmean_a
            df['dispabsmean'] = dispabsmean_a
            df['dispproba'] = dispproba_a
            df['dispabsproba'] = dispabsproba_a
            df['Fmean'] = Fmean_a
            df['Fabsmean'] = Fabsmean_a
            df['Fproba'] = Fproba_a
            df['Fabsproba'] = Fabsproba_a
            df.to_excel(saving_path+'data.xlsx')


        #Looping on HSP_ER
        if do_parallel :
            Parallel(ncores)(delayed(HSP1_calc)(HSP1) for HSP1 in HSP_ER)
        else :
            for HSP1 in HSP_ER :
                HSP1_calc(HSP1)


        #End loops Nt
        Nt_toc = time.perf_counter() #End of loop for Number of tango
        print(' took '+str(int((Nt_toc-Nt_tic)*100)/100)+' seconds')


    #Ending code
    toc = time.perf_counter()
    print('\nCode lasted '+str(int((toc-tic)*100)/100)+' seconds')
    print('*****End script*****')

# main
# =============================================================================
```

Python code providing the probabilities of biniding.

```
"""
Created on Sat Sep  2 15:08:44 2023

@authors: Lancelot Pincet, Frédéric Pincet

Gives the probabilities for a procollagen site to be bound to TANGO1

Case with :
pH 7.5 in ER, 6.5 in Golgi
10nM HSP47 in ERGIC
pH gradient, HSP47 gradient
variable HSP47 in ER
variable number of Tango1 copies per collagen
"""

#imports
import numpy as np


# =============================================================================
# User inputs

#files
path = r'' #Path where to save files

#Loops
#Ntango = [10,30,60,100,300,600] #Number of tango1 molecules
Nt = 40 #Number of tango1 molecules
HSP_ER = [1,5,10,20,30,40,50,100,500] #HSP47 concentration in ER [µM]

#Collagen definition
coll_max = 26 #maximum number of collagen sites
coll_min = 2 #minimum number of collagen sites
coll_d = 10 #Distance between two pairs of collagen sites [nm]

#Gradient on pH
pH_start = 0 #position start of gradient of pH [nm]
pH_end = 10 #position end of gradient of pH [nm]

#HSP47 concentrations
HSP_start = 0 #position start of gradient of HSP47 [nm]
HSP_end = 10 #position end of gradient of HSP47 [nm]
C_HSP2 = 1e-9 #HSP47 concentration in ERGIC [M]
Nconc = 161 #nb of considered HSP47 concentrations in ER


#Tango1 polymer description
pers = 1.4 #persistence length in nm = 4 residues
Monom = 264 #nb of monomers = 1056 residues / 4 residues
Avogadro = 6.02e-1 #in molec.nm^-3.M^-1

#Affinities
KS = 0.26e-6 #Affinity of HSP47 with SH3 [M]
KC_ERGIC = 6.23e-6 #Affinity of HSP47 with collagen site in ERGIC [M]
KC_ER = 0.74e-6 #Affinity of HSP47 with collagen site in ER [M]

# User inputs
# =============================================================================


#Inializing collagen
xpos = (np.arange(31)-15)*10 #Collagen site positions

#Initializing Tango1
cont = Monom * pers #contour length of Tango1 [nm]
sigma = np.sqrt(Monom*pers**2) #standard deviation
ppos = (3**(3/2)*np.exp(-3*xpos**2/2/sigma**2))/(2*np.pi*sigma**2)**(3/2) #density probability of SH3
alpha = abs(xpos/cont)
force = -np.sign(xpos)*4.14/pers*(1/(4*(1-alpha))-0.25+alpha-0.5164228*alpha**2-2.737418*alpha**3+16.07497*alpha**4-38.87607*alpha**5+39.49944*alpha**6-14.17718*alpha**7) #Force of polymer with end distance = alpha*contour_length
xpos_accurate = np.arange(201)-100
ppos_accurate = (3**(3/2)*np.exp(-3*xpos_accurate**2/2/sigma**2))/(2*np.pi*sigma**2)**(3/2) #density probability of SH3
ppos_norm = ppos_accurate/np.sum(ppos_accurate)

#Affinity of HSP47 with collagen site [M]
KC = np.empty(np.shape(xpos))
mask_pH1 = xpos < pH_start
mask_pH2 = (xpos >= pH_start) * (xpos <= pH_end)
mask_pH3 = xpos > pH_end
KC[mask_pH1] = KC_ER
KC[mask_pH2] = KC_ER + (KC_ERGIC - KC_ER)*(xpos[mask_pH2] - pH_start)/(pH_end-pH_start)
KC[mask_pH3] =KC_ERGIC

#Variable initializations
probaS = np.empty(np.shape(xpos)) #Probability that an SH3-like domain is bound to HSP47
probaC = np.empty(np.shape(xpos)) #Probability that a collagen binding-site is bound to HSP47

# main
CHSPe = []
F_mean = []
f_bound = []
Nb_bonds = []

for NC1 in range(Nconc) :
    #Start C_HSP1
    C_HSP1 = 10**((NC1-60)/20-6) #HSP1 in µM C_HSP1 in M (ER)
    CHSPe += [10**((NC1-60)/20)]
    HSP2 = C_HSP2 * 1e+6 #HSP2 in µM C_HSP2 in M (ERGIC)

    #HSP47 concentrations
    C_HSP = np.empty(np.shape(xpos))
    mask_HSP1 = xpos < HSP_start
    mask_HSP2 = (xpos >= HSP_start) * (xpos <= HSP_end)
    mask_HSP3 = xpos > HSP_end
    C_HSP[mask_HSP1] = C_HSP1
    C_HSP[mask_HSP2] = C_HSP1 + (C_HSP2 - C_HSP1)*(xpos[mask_HSP2] - HSP_start)/(HSP_end-HSP_start)
    C_HSP[mask_HSP3] = C_HSP2

    #Probabilities
    probaS = 1/(1+KS/C_HSP)
    probaC = 1/(1+KC/C_HSP)
    CbT = probaS*Nt*ppos/Avogadro #local concentration of HSP47-bound SH3-like domains
    CuT = (1-probaS)*Nt*ppos/Avogadro #local concentration of HSP47 not bound to SH3-like domains
    #p_tot = (probaC/(1+KC/CuT)+(1-probaC)/(1+KC/CbT)).astype(np.float32) #probability to have a collagen - Tango bond at each collagen site
    p_tot = (1/(1+KC/CbT+KS/CuT)).astype(np.float32) #probability to have a collagen - Tango bond at each collagen site
    f_bound += [(np.sum(p_tot)*2)/Nt] #the 2 comes from the 2 sites located at the same position
    p_norm = p_tot/np.sum(2*p_tot) #because of the 2 sites
    F_mean += [np.sum(force*p_norm)] #because of the 2 sites
    Nb_bonds += [np.sum(p_tot)]


#50µM HSP1
C_HSP1 = 5e-5 #HSP1 in µM C_HSP1 in M (ER)
#HSP47 concentrations
C_HSP = np.empty(np.shape(xpos))
mask_HSP1 = xpos < HSP_start
mask_HSP2 = (xpos >= HSP_start) * (xpos <= HSP_end)
mask_HSP3 = xpos > HSP_end
C_HSP[mask_HSP1] = C_HSP1
C_HSP[mask_HSP2] = C_HSP1 + (C_HSP2 - C_HSP1)*(xpos[mask_HSP2] - HSP_start)/(HSP_end-HSP_start)
C_HSP[mask_HSP3] = C_HSP2

#Probabilities
probaS = 1/(1+KS/C_HSP)
probaC = 1/(1+KC/C_HSP)
CbT = probaS*Nt*ppos/Avogadro #local concentration of HSP47-bound SH3-like domains
CuT = (1-probaS)*Nt*ppos/Avogadro #local concentration of HSP47 not bound to SH3-like domains
p_tot_50 = (1/(1+KC/CbT+KS/CuT)).astype(np.float32) #probability to have a collagen - Tango bond at each collagen site

#1µM HSP1
C_HSP1 = 1e-6 #HSP1 in µM C_HSP1 in M (ER)
#HSP47 concentrations
C_HSP = np.empty(np.shape(xpos))
mask_HSP1 = xpos < HSP_start
mask_HSP2 = (xpos >= HSP_start) * (xpos <= HSP_end)
mask_HSP3 = xpos > HSP_end
C_HSP[mask_HSP1] = C_HSP1
C_HSP[mask_HSP2] = C_HSP1 + (C_HSP2 - C_HSP1)*(xpos[mask_HSP2] - HSP_start)/(HSP_end-HSP_start)
C_HSP[mask_HSP3] = C_HSP2

#Probabilities
probaS = 1/(1+KS/C_HSP)
probaC = 1/(1+KC/C_HSP)
CbT = probaS*Nt*ppos/Avogadro #local concentration of HSP47-bound SH3-like domains
CuT = (1-probaS)*Nt*ppos/Avogadro #local concentration of HSP47 not bound to SH3-like domains
p_tot_1 = (1/(1+KC/CbT+KS/CuT)).astype(np.float32) #probability to have a collagen - Tango bond at each collagen site


# main
# =============================================================================
```

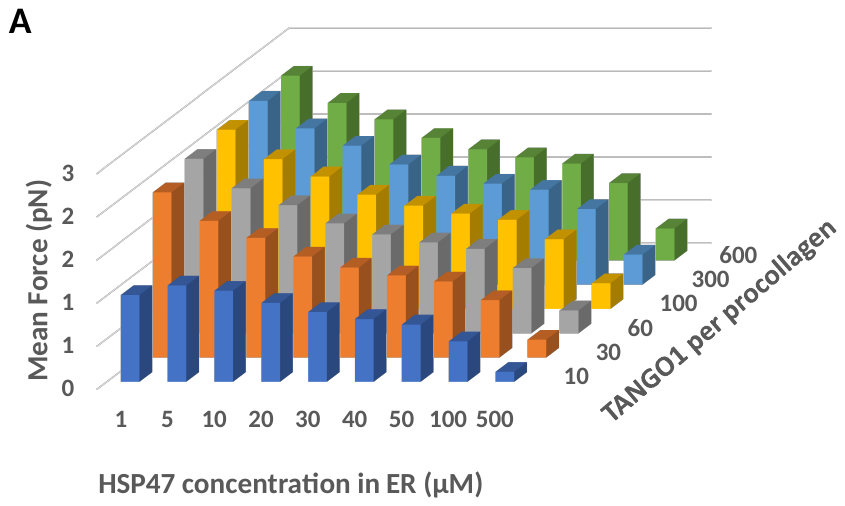

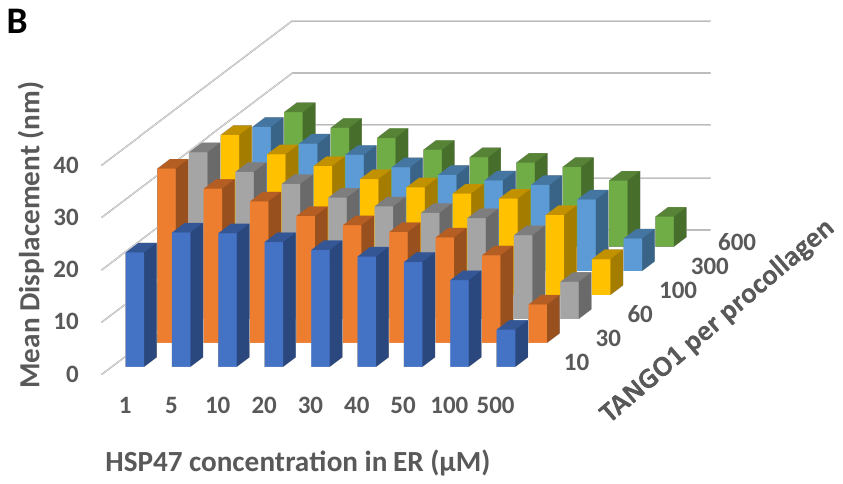

Fig. S1. Equilibrium: force and displacement for a stepwise drop of HSP47 to 1 nM at the base of the tunnel and an affinity (KD) varying linearly over 60 nm from 0.74 µM in the ER to 2.25 µM in the ERGIC over, corresponding to a pH gradient. A. Mean force on procollagen when several TANGO1 (10 to 40, right axis) are acting at equilibrium and for HSP47 concentrations varying from 1 µM to 100 µM in the ER lumen (front axis). B. Mean procollagen displacement (step) to dissipate the force exerted by Tango1 on procollagen.

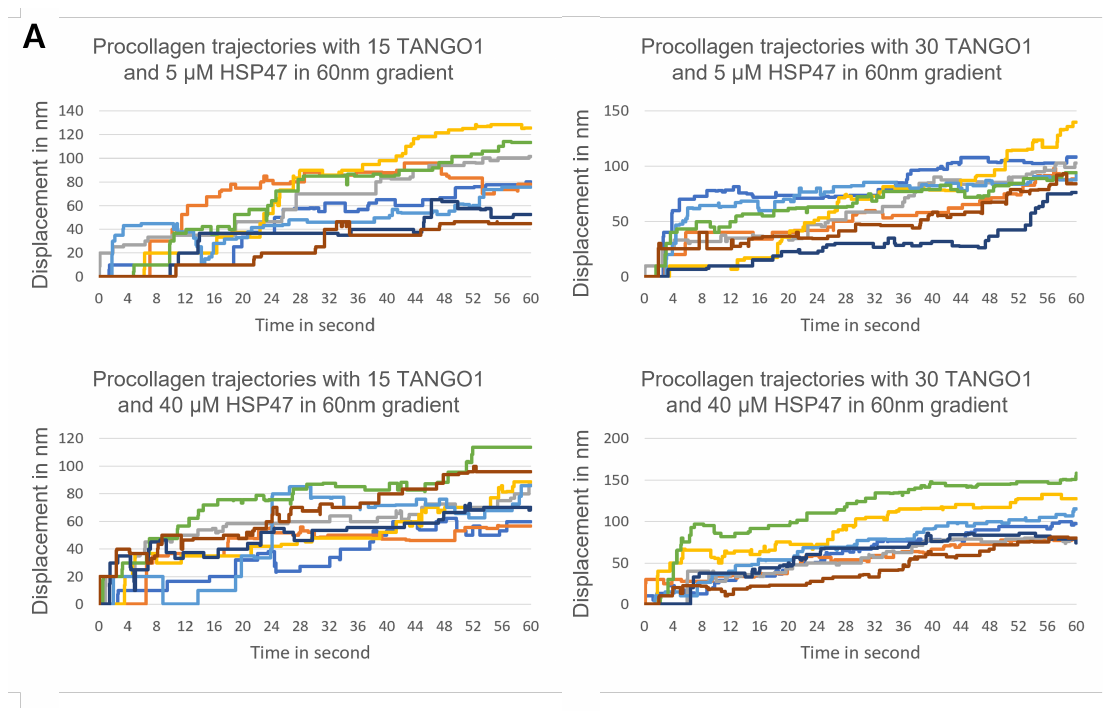

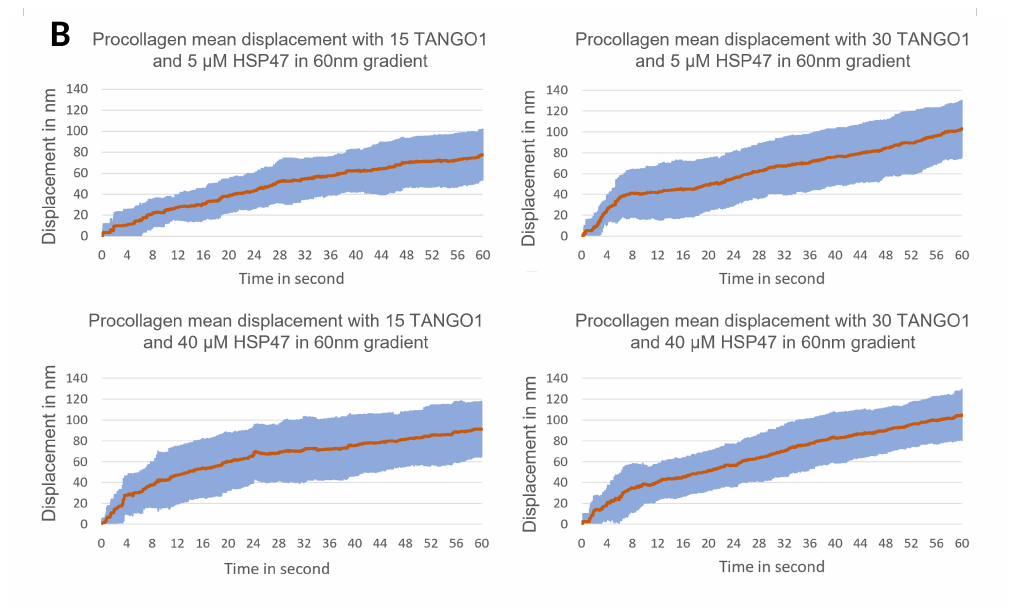

**Fig. S2.** Movement of unconstrained procollagen for a stepwise drop of HSP47 to 100 nM at the base of the tunnel and an affinity (K_D_) varying linearly from 0.74 µM at the base of the tunnel, $x=0$, to 2.25 µM at $x=60 nm$ in the tunnel, corresponding to a pH gradient. A. Examples of trajectories (displacement of procollagen) in time for 8 simulations per conditions. B. Mean of the displacements over 15 trajectories. The blue shaded area corresponds to the standard deviation. In A and B, left (resp. right) panels are for 15 (resp. 30) TANGO1 and Top (resp. Bottom) panels are for 5 µM (resp. 40 µM) HSP47 in the ER lumen.

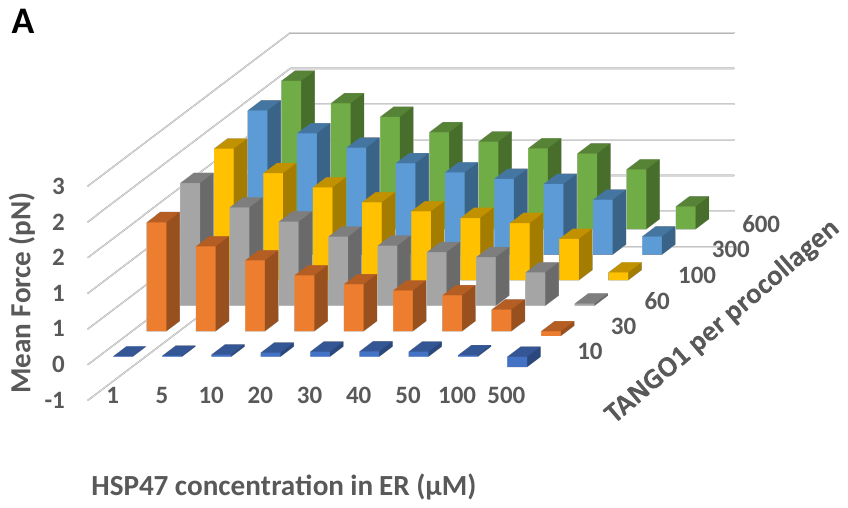

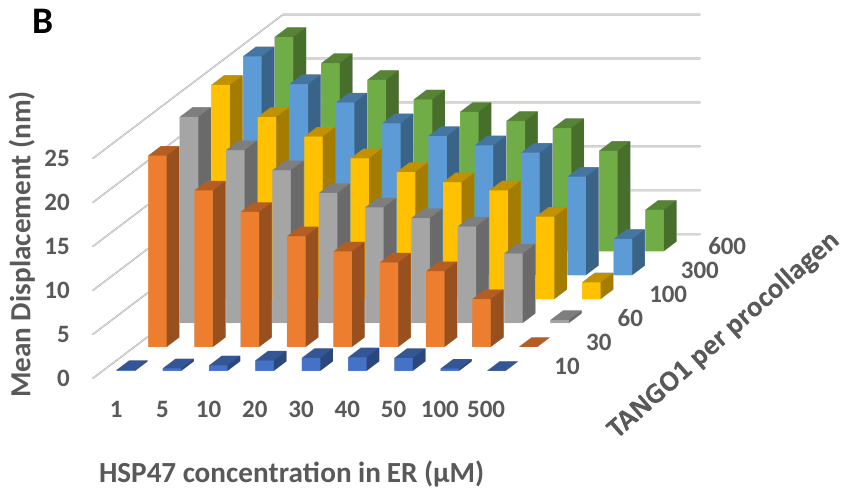

Fig. S3. Equilibrium: force and displacement for a HSP47 concentration varying linearly ovr 60 nm from 1 nM in the ER and 1 nM in th ERGIC and an affinity (KD) varying linearly over 60 nm from 0.74 µM in the ER to 2.25 µM in the ERGIC over, corresponding to a pH gradient. A. Mean force on procollagen when several TANGO1 (10 to 40, right axis) are acting at equilibrium and for HSP47 concentrations varying from 1 µM to 100 µM in the ER lumen (front axis). B. Mean procollagen displacement (step) to dissipate the force exerted by Tango1 on procollagen.

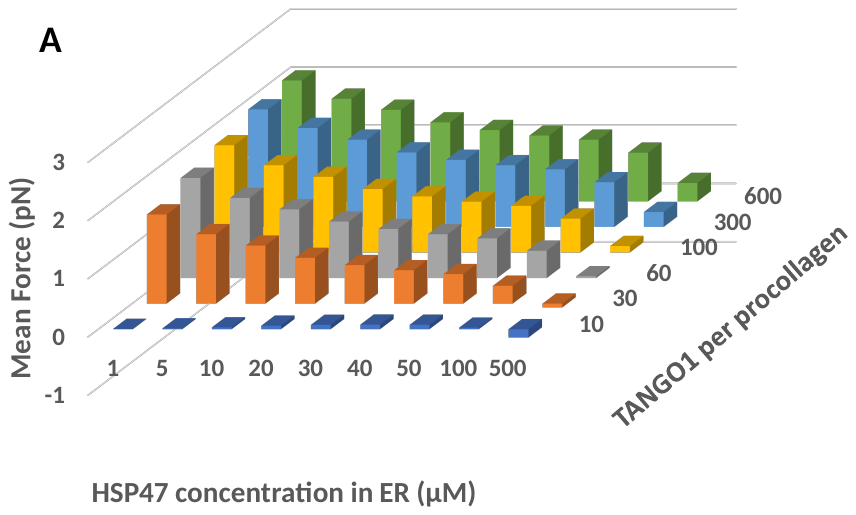

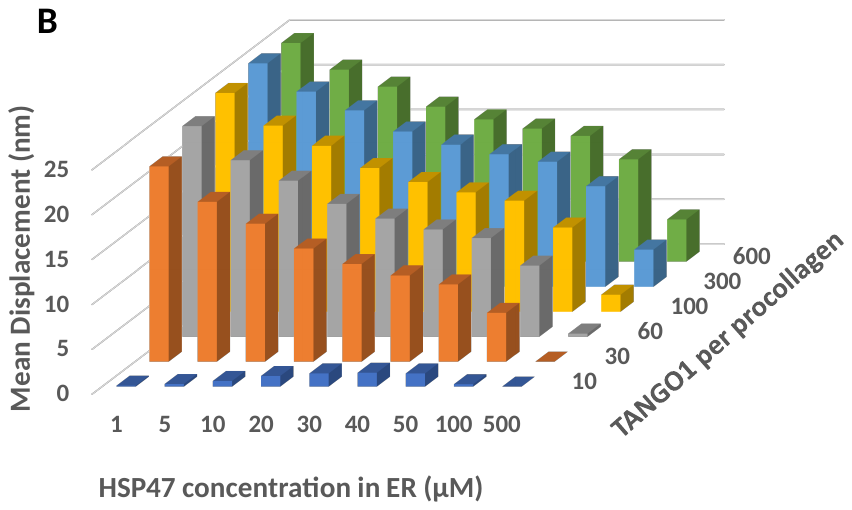

Fig. S4. Equilibrium: force and displacement for an HSP47 concentration varying linearly ovr 60 nm from 1 nM in the ER and 1 nM in th ERGIC and a stepwise variation of the pH from 7.5 to 6.5. A. Mean force on procollagen when several TANGO1 (10 to 40, right axis) are acting at equilibrium and for HSP47 concentrations varying from 1 µM to 100 µM in the ER lumen (front axis). B. Mean procollagen displacement (step) to dissipate the force exerted by Tango1 on procollagen.

1.	Ishikawa Y, Ito S, Nagata K, Sakai LY, & Bachinger HP (2016) Intracellular mechanisms of molecular recognition and sorting for transport of large extracellular matrix molecules. *Proc Natl Acad Sci U S A* 113(41):E6036-E6044.

2.	Macdonald JR & Bachinger HP (2001) HSP47 binds cooperatively to triple helical type I collagen but has little effect on the thermal stability or rate of refolding. *J Biol Chem* 276(27):25399-25403.

3.	Stirnemann G, Giganti D, Fernandez JM, & Berne BJ (2013) Elasticity, structure, and relaxation of extended proteins under force. *Proc Natl Acad Sci U S A* 110(10):3847-3852.

4.	Itzhak DN, Tyanova S, Cox J, & Borner GH (2016) Global, quantitative and dynamic mapping of protein subcellular localization. *Elife* 5.

5.	Andrews SS (2005) Serial rebinding of ligands to clustered receptors as exemplified by bacterial chemotaxis. *Phys Biol* 2(2):111-122.

6.	Broersma S (1981) Viscous Force and Torque Constants for a Cylinder. *Journal of Chemical Physics* 74(12):6989-6990.

7.	Oecal S*, et al.* (2016) The pH-dependent Client Release from the Collagen-specific Chaperone HSP47 Is Triggered by a Tandem Histidine Pair. *J Biol Chem* 291(24):12612-12626.

8.	Natsume T, Koide T, Yokota S, Hirayoshi K, & Nagata K (1994) Interactions between collagen-binding stress protein HSP47 and collagen. Analysis of kinetic parameters by surface plasmon resonance biosensor. *J Biol Chem* 269(49):31224-31228.
